# Ancient genomics reveals a genetic continuum with dual structure in the Classic Copan

**DOI:** 10.1101/2024.05.17.594672

**Authors:** Madeleine Murray, Seiichi Nakamura, Melvin Fuentes, Masahiro Ogawa, Lara M. Cassidy, Takashi Gakuhari, Shigeki Nakagome

## Abstract

Copan was a major capital at the southeasternmost of the Maya civilization, serving as a crossroads connecting Central and South America. In 426/427 CE, the city witnessed the establishment of a royal dynasty, which endured for approximately 400 years. Despite extensive historical and archaeological records, there remains limited information regarding the genetic origins of people who resided in Copan. Here, we present seven Classic Copan genomes including an enigmatic dynastic ruler and his sacrificed companion. Our analysis identifies a high genetic affinity of the Classic Copan with present-day Maya. Both groups exhibit an admixture of Mesoamerican and South American Farmer (SAF) ancestry, with a shift in a dominant component from SAF to Mesoamerican. At the individual level, commoners show a more prominent SAF ancestry, while the royal individual demonstrates nearly equal contributions from two ancestors. These results highlight a dual ancestral structure in this geographic extreme of the Classic Maya state, uncovering a cline of Mesoamerican ancestry across individuals.

## Introduction

Copan was one of the most important centers across the Classic Maya states, distinguished by its well-preserved sculptures and the hieroglyphic stairway, which features the longest Maya text inscription found in the Pre-Columbian Americas. The earliest human occupation at Copan goes back to the Early Pre-Classic, or possibly even the Late Archaic period. This period is marked by the emergence of small farming communities, primarily relying on maize-based food production, in the Copan Valley (*1*, *2*), located in western Honduras near the border of Guatemala. Monumental construction began in this region during the Early Classic period, approximately between 300-400 CE. Copan’s royal dynasty was then founded in 426/427 CE, according to the current results of epigraphic decipherment (*3*). The dynastic succession saw 16 rulers over the ancient city and its polity. Copan thrived as a political, economic, and ceremonial hub, encompassed by elite and commoner residential structures. While the population is estimated to have peaked at around 30,000 inhabitants (*4*), the political regime ultimately collapsed *circa* 820 CE at the end of the Late Classic period (*5–7*).

A long-standing hypothesis on the formation of the Copan kingdom proposes the presence of an enclave of foreign Maya within a territory inhabited by non-Maya people (*7–9*). Individuals from a widely dispersed Maya elite network migrated potentially from the central Petén region of Guatemala, especially around Tikal area, to the region, intermarried with local Maya residents, and settled in the area until their passing. Archaeological evidence points to extensive trade and political connections between Copan and other Mesoamerican cities, such as Tikal, Kaminaljuyú in the Valley of Guatemala, and Teotihuacán in the Valley of Mexico (*2*, *5*, *7*). Inscriptions from Copan reveal that the first dynastic founder, K’inich Yax K’uk’ Mo’, was an outsider who assumed power in 426 CE and arrived at Copan in 427 CE (*3*, *10*, *11*). Isotopic analysis with tooth enamel provides further support on the archaeological and epigraphic data by uncovering significant variation in the places of origins among human remains discovered at the Copan sites (*12*). Notably, this variation is more pronounced among elites, including the first and subsequent kings, than among commoners (*13*, *14*). The chemistry of tooth enamel reflects the foods and liquids consumed by the first generation immigrants during their infancy and early childhood (*15*). Since those elites buried at the Copan sites are typically adults (*14*), this suggests that these elites grew up outside of Copan before relocating to the city. However, distinguishing foreigner status through visible attributes, such as dental decoration, artificial head shaping, and osteological characters, is challenging (*16*, *17*); whether the social and cultural assimilation process was accompanied by genetic exchanges remains elusive.

Here, we report seven newly sequenced ancient Copan genomes, all originating from the Classic Maya period (Table 1). To our knowledge, this dataset represents the first collection of ancient genomes from the Classic Maya state, including those of an enigmatic dynastic ruler who was buried outside of the Acropolis (CpM13) and an individual discovered in close proximity to the ruler (CpM12), likely associated with a sacrificial burial, as well as commoners (*14*). By integrating these Copan genomes with a broader ancient and modern genomic dataset spanning North-to-South Americas and Siberia, our study aims to enhance our understanding of the genetic ancestry of the Classic Copan individuals, shedding light on the dynamics of population movement and social networks at the southeastern edge of the Maya region.

**Table 1.**
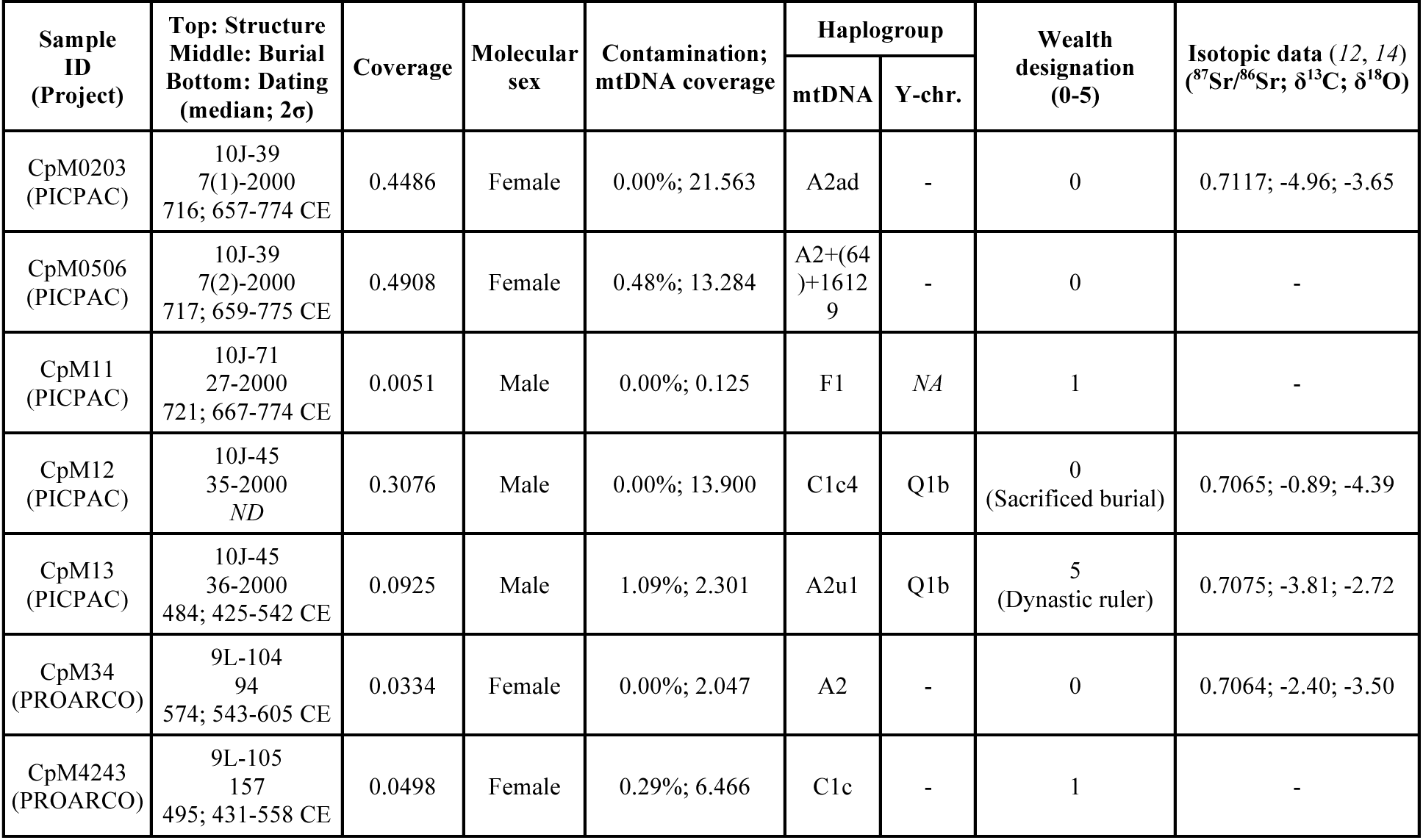
Summary of Classic Copan genomes and their associated profiles.

## Results

### Ancient genomes from the Classic Copan

Our initial screening focused on 16 ancient skeletal remains excavated from the sites of Copan, as part of two projects (PICPAC and PROARCO), both led by S. Nakamura (see note S1). More than 1% of endogenous human DNA were preserved in seven of these samples (Table 1), which were then further shotgun-sequenced to higher coverage, ranging from 0.033× to 0.491×. Those seven individuals are associated with five different structures (PICPAC: 10J-39, 10J-45, and 10J-71; PROARCO: 9L-104, and 9L-105). Radiocarbon dating conducted on collagen samples extracted from the corresponding bones reveals that all samples align with the Classic Maya period, yet they are split into two temporal groups: either the Early-to-Middle or the Middle-to-Late Classic Maya phase. A wealth designation, measured from various markers including the structure where they were buried and the number, size, and complexity of accompanied burial offerings, represents roughly the social standing of a buried individual, with “0” or “5” being the lowest or the highest. The burial form, contexts, and contents, marked by the vaulted stone chamber with two stone burial slabs, as well as the presence of exquisite jade offerings inside and outside of the chamber provide compelling evidence that CpM13 held a position as one of Copan’s dynastic rulers (*14*, *18*). Notably, this individual was accompanied with two jade pectorals. One of these pectorals features intricate engravings of mat motifs, symbolizing the rulership and divine power of the buried individual (*19*). The second pectoral displays a carved motif of the Pax god, signifying the military prowess of the entombed individual. Adjacent to the burial site of CpM13 within the same structure (10J-45), CpM12 was found beneath the wall of the basement at the funerary temple constructed for CpM13, without accompanying any offering; CpM12 is then identified as a sacrificial companion (*14*, *18*).

We confirm that all newly sequenced genomes show postmortem damage patterns (fig. S1) and demonstrate a low level of modern human contamination (<1.09%) (Table 1). Our kinship analysis supports potential second-degree relationships between CpM11 and CpM34 or between CpM11 and CpM4243 (fig. S2). Except for CpM11 that was inferred for a mitochondrial haplogroup of the F1 clade from a low coverage of mitochondrial genome, the haplogroups fall within the A2 or C1 clades, both classified as Pan-American mtDNA haplogroups and commonly observed in present-day populations throughout the Americas (*20–22*). Molecular sexing identifies three males, including the sacrificed victim (CpM12), previously considered female based on morphological characteristics (*14*, *23*). The two males, CpM13 and CpM12, belong to Q1b, the most prevalent haplogroup in Native American populations (*24*), while the other male, CpM11, has insufficient sequence coverage for determining the Y chromosome haplogroup. To contextualise our data within the broader landscape of American demography, we integrated the ancient Copan genomes with genomic data from previously published ancient individuals (table S1 and fig. S3) as well as present-day populations.

### Population structure of the Classic Copan

We explored the genome-wide autosomal affinities of our Classic Copan individuals to the other ancient individuals representative of North, Central, and South America, along with present-day populations, using principal components analysis (PCA). By projecting ancient individuals onto the genetic variation of present-day populations in the Human Origin Array dataset from the Americas (Fig. 1A and fig. S4), we observe that the Copan individuals form a distinct cluster, closely located to clusters of Late Archaic genomes from Belize (*25*)and pre-Hispanic genomes from Panama (*26*). Within the Copan, some individuals exhibit a slight shift toward present-day Maya individuals, indicating a potential genetic link between ancient and modern individuals associated with Maya culture. Additionally, there is a cline defined by ancient Peruvian and Bolivian individuals (*27*, *28*), with a clear direction toward present-day Bolivian individuals. These patterns remain consistently observable across different sets of present-day reference populations (fig. S4).

**Figure 1.**
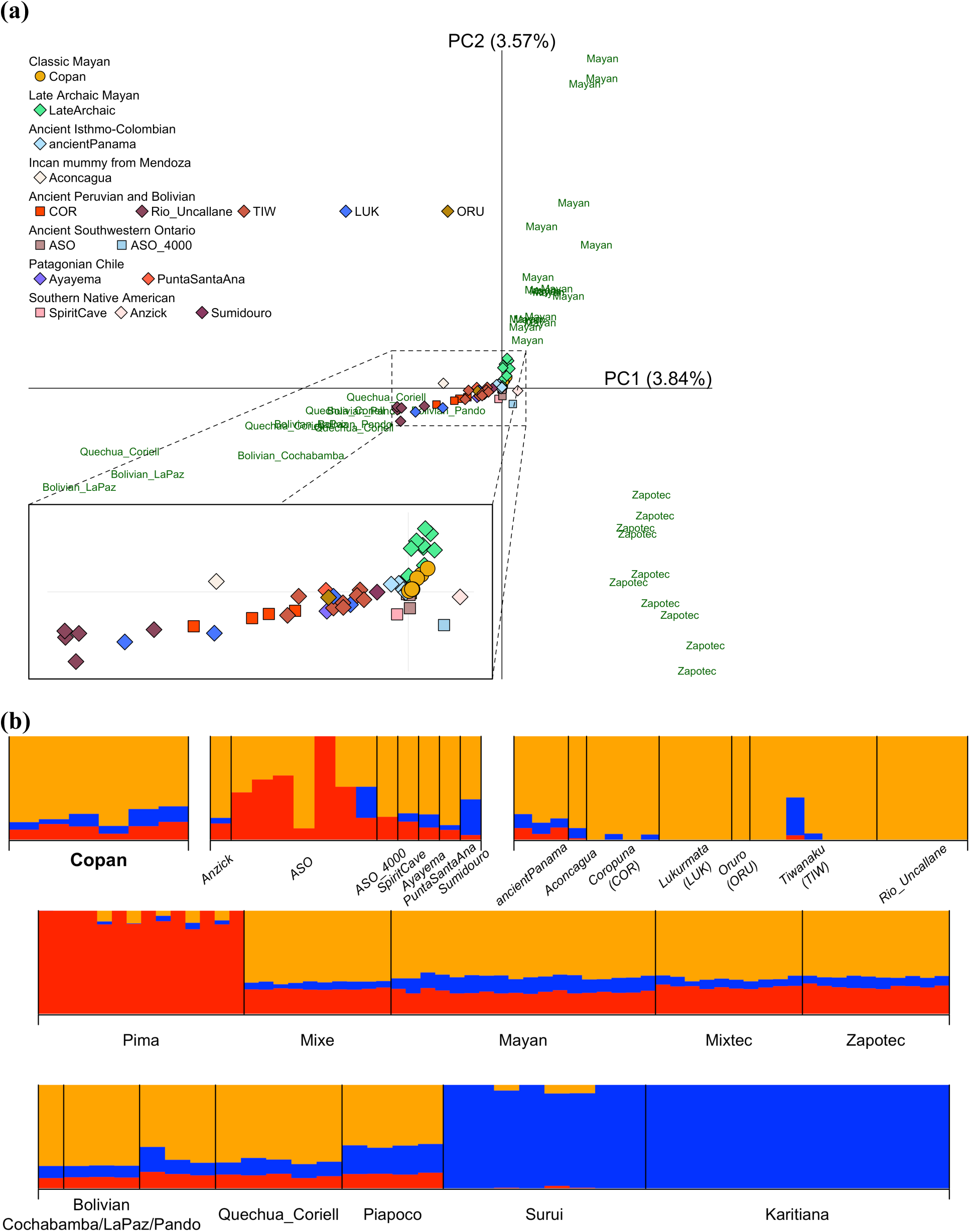
Genomic profiles of the Classic Copan individuals in the context of ancient and present-day American populations. **(a)** Principal components analysis (PCA) visualizing Classic Copan individuals and other ancient individuals from the Americas (presented as colored symbols) projected onto present-day American populations (shown by dark green with their own population labels). The same set of ancient individuals is projected onto different sets of present-day American populations in fig. S4. **(b)** ADMIXTURE analysis with ancient (bold and italic for the Classic Copan and italic for the others) and present-day American individuals (*K* = 3; results from *K* = 2 to *K* = 7 are presented in fig. S6). Within the Classic Copan, six individuals are aligned in the following order: CpM0203, CpM0506, CpM12, CpM13, CpM34, and CpM4243. The ancient individuals includes following individuals or populations: Anzick (a late Pleistocene individual from a Clovis burial site in western Montana), ASO (an ancient Southwestern Ontario population), ASO_4000 (an ASO individual dated to 4,267 ± 22 calBP), SpiritCave (a 10,700-years-old individual from Nevada), Ayayema and PuntaSantaAna (a 5,100-years-old and a 7,200-years-old individuals from Patagonia in Chile), Sumidouro5 (an ancient Amazonian individual from Brazil with a date around 10,000 years ago), ancientPanama (pre-Hispanic genomes from Panama), Aconcagua (a 500-years-old frozen mummy discovered in Argentina, who are considered to have been brought from the Central Coast of Peru), Coropuna (ancient southern Peruvian individuals associated with either the Wari or Inca culture; the population is abbreviated as COR), Lukurmata (ancient Bolivian individuals from the Lake Titicaca basin spanning the period from before the Tiwanaku civilisation to after its abandon; LUK), Oruro (an ancient Bolivian individual discovered from a site approximately 200 km away from a site where LUK individuals were excavated, with a calibrated date of 1,410 CE corresponding to the historic period between the post-Tiwanaku and pe-Inca civilisations; ORU), Tiwanaku (ancient individuals from the ritual core of the Tiwanaku site in the Bolivian Lake Titicaca region; TIW), and Rio_Uncallane (ancient sedentary farmers from the Lake Titicaca region of Peru, which are dated to approximately 1,800 BP).

ADMIXTURE analysis performed for *K* = 4, chosen as the point just before cross-validation errors significantly increase (fig. S5), reveals three distinct ancestors characterising Pima (red), an indigenous population from Mexico, and two Amazonian populations: Surui (magenta) and Karitiana (blue) (Fig. 1B). These populations clearly stand out in our PCA (fig. S4A), supporting their genetic uniqueness across American populations. The other component coloured by orange represents an ancestor of Rio Uncallane, sedentary agriculturalists dating back 1,800 years in the Lake Titicaca region of Peru (*28*). This orange component is dominant for ancient Peruvian and Bolivian individuals, who are associated with Tiwanaku, Wari, or Inca culture (*27*), as well as a 500-year-old frozen mummy found in Argentina (*29*). All four ancestral components are present in the Copan individuals, with the orange and red components comprising over 70% of their genetic ancestry. This pattern observed in the Copan individuals is also visible in present-day populations from Central America or Bolivia, as well as in most of the other ancient individuals included in this analysis.

### Genetic affinities to the Classic Copan

To investigate genetic relationships between the Classic Copan and the populations from Siberia and the Americas (including 53 ancient and 12 present-day populations), we computed *f*_4_-statistics with the form of *f*_4_(Mbuti, *X*; *Y*, Classic Copan). This analysis iterates through each of these 65 populations as *X*, with the remaining populations designated as *Y*. We then ranked the populations based on the frequency with which a population exhibits a greater genetic affinity to the Copan compared to each of the 64 other populations, with a *Z*-score > 3.0 (Figure 2). This comprehensive analysis clearly demonstrates that strong genetic affinities to the Copan are observable among the populations from Central America. In particular, the present-day Maya emerges as the top population, which is further supported by the genetic affinities observed in five out of six Classic Copan individuals when analysed at the individual level (fig. S7 and table S2). These results suggest a certain level of genetic continuity between the Classic Maya period and the present. CpM34 stands out as the only Copan individual who does not support genetic affinity with the Maya population. Instead, the top-ranked population is LUK, a group of ancient Bolivian individuals from both pre- and post-Tiwanaku civilization (*27*), although it is worth noting that the low coverage of data may have influenced these results.

**Figure 2.**
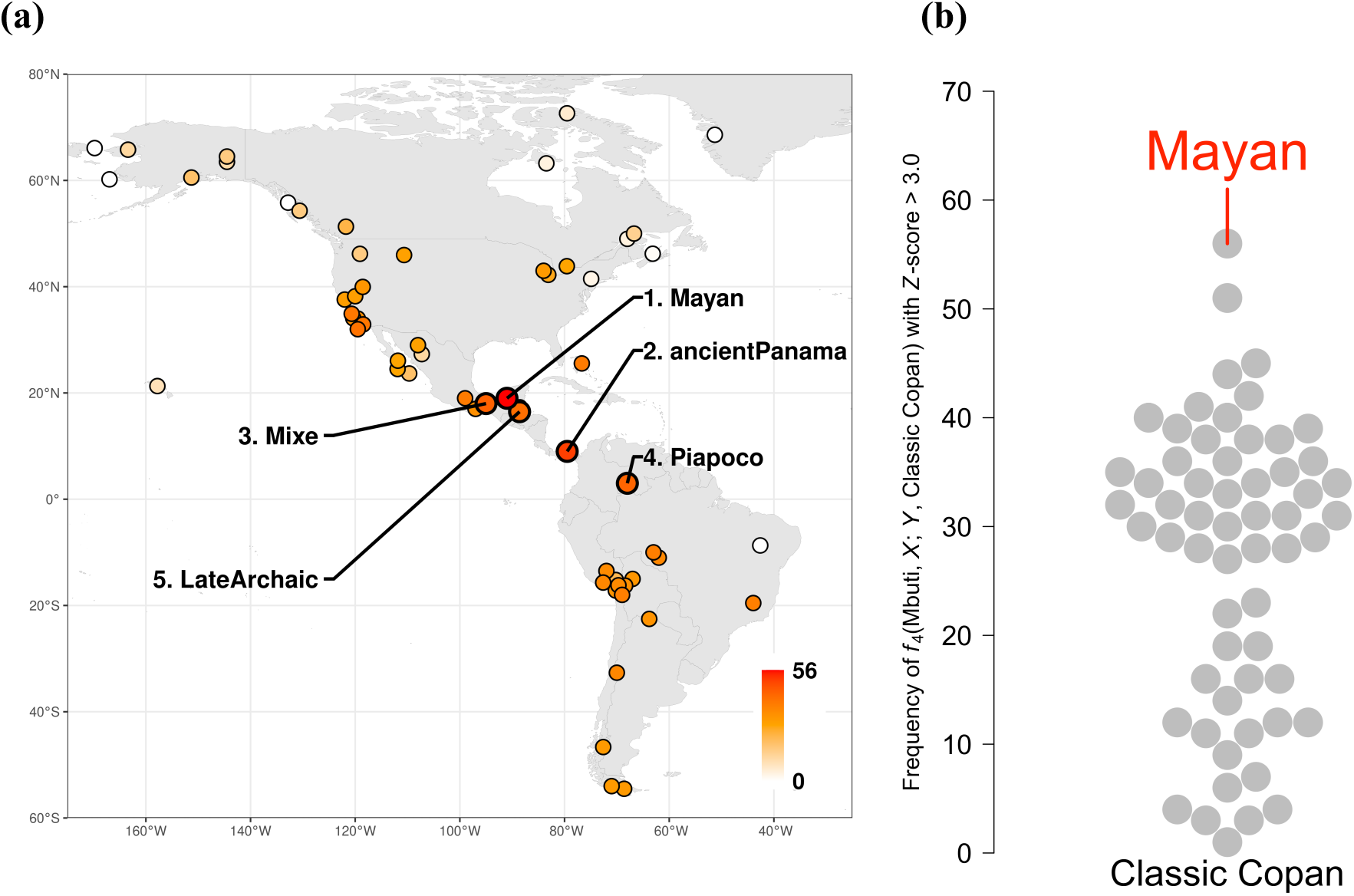
Ancient and present-day populations ranked by significant genetic affinities to the Classic Copan in the form of *f*_4_(Mbuti, *X*; *Y*, Classic Copan). **(a)** A geographical map illustrates a population *X* who exhibits an excess genetic affinity to the Classic Copan compared to a population *Y*, with a *Z*-score > 3.0. The color gradation reflects the frequency with which a population stands out in the *f*_4_-tests. The top five populations are labelled on the map. Three populations from Siberia, AG2, MA1 and Tlingit, are included in the tests, but are not shown on the map, all of which are positioned with lower rankings in the tests. **(b)** The plots represent the frequencies and rankings of all populations mapped in (a). The present-day Maya population emerges as the most prevalent outlier. Results of frequencies and rankings at the individual level are presented in fig. S7 and table S2.

We then explore whether the top populations (*i*.e., Mayan or LUK) alone could fully account for the genetic ancestry of the Copan. There are certain instances in *f*_4_(Mbuti, *Top_population*; *Y*, Classic Copan/CpM), where the genetic affinity between the top populations and the Copan did not reach statistical significance. This could be attributed to factors such as insufficient statistical power (*e*.*g*., a small sample size or a small number of SNPs) or a scenario where population *Y* also shares genetic ancestry with the Copan. To test the latter, we employed *f*_4_-tests with the form of *f*_4_(Mbuti, *Y*; *Top_population*, Classic Copan/CpM), where *Top_population* is substituted with *Y* in the previous form of *f*_4_-tests. However, none of the populations tested as *Y* in *f*_4_(Mbuti, *Top_population*; *Y*, Classic Copan/CpM), where a *Z*-score was below the statistical significance, show genetic affinity to the Copan relative to the top populations (fig. S8). These results suggest that the top populations adequately represent the genetic ancestors characterising the Classic Copan.

### Genetic origins of present-day Maya

The present-day Maya are widely considered genetically as a Mesoamerican population. In fact, the Maya derive approximately 80% of their genetic ancestry from Mixe, a well-known population that broadly represents Mesoamerican ancestors (*29*). Consistent with these earlier observations, our *f*_4_-tests, expressed as *f*_4_(Mbuti, *X*; *Y*, Mayan), rank Mixe as the top population *X* (Figure 3a). Meanwhile, there are also cases where populations, acting as *Y* in *f*_4_(Mbuti, Mixe; *Y*, Mayan), do not lend support to the genetic affinity between Mixe and Mayan, which may suggest a dual-ancestral structure of the Maya population. Indeed, additional *f*_4_-tests with the form of *f*_4_(Mbuti, *X*; Mixe, Maya), where *X* represents populations with *Z* < 3.0 in *f*_4_(Mbuti, Mixe; *Y*, Mayan), reveal that six populations exhibit a greater affinity to the Maya relative to Mixe (fig. S9).

**Figure 3.**
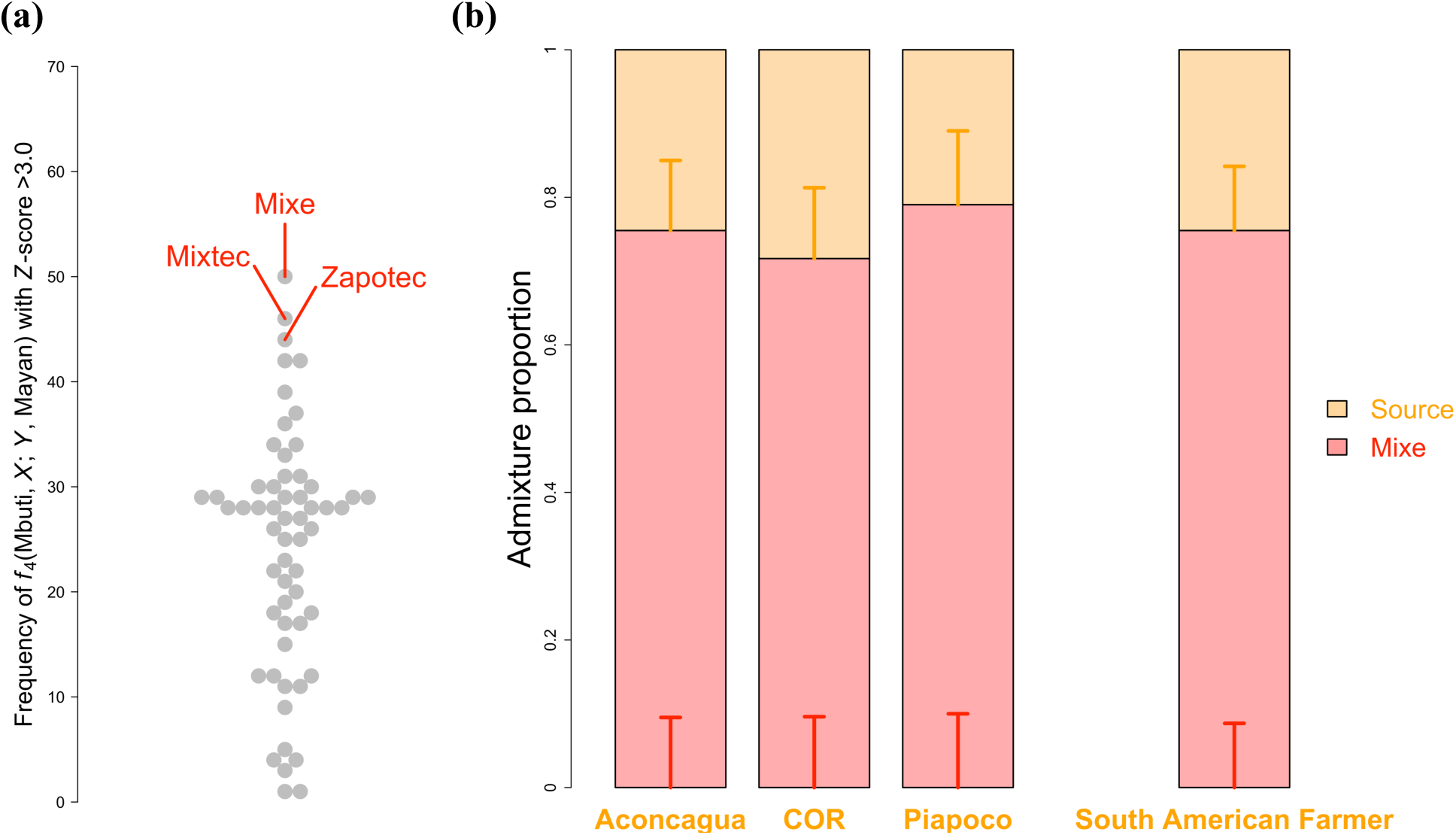
Genetic ancestry of the present-day Maya. **(a)** Genetic affinity of ancient and modern populations from Siberia and the Americas to the Maya population is ranked by the number of times each tested population stands out as a population *X* in the form of *f*_4_(Mbuti, *X*; *Y*, Maya) with a *Z*-score >3. The highest occurrence of a population *X* is Mixe, followed by Mixtec and Zapotec. **(b)** Genetic ancestry of the Maya population is modelled with a two-way admixture of Mixe and each of three potential sources that all exhibit an extra genetic affinity to the Maya compared with Mixe (see fig. S9). Vertical bars represent ±1 SE estimated by *qpAdm*. Given shared genetic ancestry between Aconcagua and COR (see Figure 2; figs. S6 and S10), these populations are grouped to represent South American Farmer (SAF) ancestry in admixture modelling. The values of admixture proportions, as well as fittings of different models, are shown in table S3.

To understand the genetic makeup of the Maya, we modelled the Maya as a two-way admixture of Mixe and each of these six potential sources using *qpAdm* (Figure 3b and table S3). The admixture models show a significantly better fit to the genetic profile of the Maya than single-ancestry models solely relying on Mixe (nested *p*-values < 0.05), when Aconcagua, COR, or Piapoco are considered as the additional ancestral sources. Admixture fractions of Mixe ancestry are consistent among those three sources: 75.5% ± 9.5% (Aconcagua), 71.7% ± 9.6% (COR), and 79.0% ± 10.0% (Piapoco). These proportions of Mixe ancestry are also in line with the previous estimates (*29*). Aconcagua, a frozen mummy from Argentina, dates back to approximately 500 years ago (*29*), which coincides with the time of southward expansion of the Inca civilization. Additionally, archaeological evidence suggests the possibility that the Aconcagua individual may have been transported from Peru (*30*). In support of this hypothesis, our analysis based on *f*_4_-statistics identifies significant genetic links between Aconcagua and COR as well as the other ancient Andean populations, including those associated with sedentary agriculture (Rio Uncallane) (*28*) or Inca and pre-Inca cultures such as Tiwanaku or Wari (TIW and LUK) (*27*) (fig. S10 and table S4). All of these ancient populations form a genetic cluster in the ADMIXTURE profiles (see the orange component in Figure 2 and fig. S6), which we define as broadly South American Farmer (SAF) ancestry.

We further confirm this dual-ancestral structure of the Maya by merging Aconcagua and COR into a single source population representing SAF ancestry in the admixture modelling (Figure 3b and table S2). An alternative model for the genetic origin of the Maya has been proposed in a previous study, suggesting that the Maya originated from admixture between Mixe and a Late Archaic Maya group from Belize (*25*). To compare the fits of two competing admixture scenarios—specifically, one involving Mixe and Late Archaic, and the other involving Mixe and SAF—we incorporated three possible ancestral sources into admixture modelling of the Maya (table S5). Contrary to the previous finding, a single ancestry of Mixe can sufficiently explain the genetic makeup of the Maya when the admixture is modelled by Mixe and Late Archaic (nested *p*-value = 0.174). However, the admixture between Mixe and SAF shows better fittings than either of the single ancestry models (*i*.*e*., Mixe-only: nested *p*-value = 0.008; SAF-only: nested *p*-value = 1.110e-16). These results strongly support the dual structure of Mesoamerican and SAF ancestry in present-day Maya populations.

### Genetic ancestry of the Classic Copan

Given the robust signals of genetic affinities between the Classic Copan and present-day Maya, we assess the applicability of the dual-ancestral structure to the genetic makeup of Copan (Figure 4a and table S6). The two-way admixture model shows a better fit to the Copan than single-ancestry models of either of the ancestors (*p*-values for the nested models < 0.05), uncovering admixture proportions of 21.0% Mesoamerican and 79.0% SAF ancestry, with a standard error of ± 8.4%. This ancestral composition in the Classic Copan highlights a significant increase in Mesoamerican ancestry between the Classic period and the present. The *f*_4_(Mbuti, *X*; Mayan, Classic Copan) tests confirm stronger genetic connections between the Copan and the populations associated with SAF ancestry compared with those in the Maya (including TIW or Rio Uncallane) (table S7). With *DATES* (*31*), we estimated that, while the variance is substantial, the admixture between Mesoamerican and SAF ancestry occurred approximately 5,000 years ago (fig. S12), the middle of the Late Archaic period in Central America.

**Figure 4.**
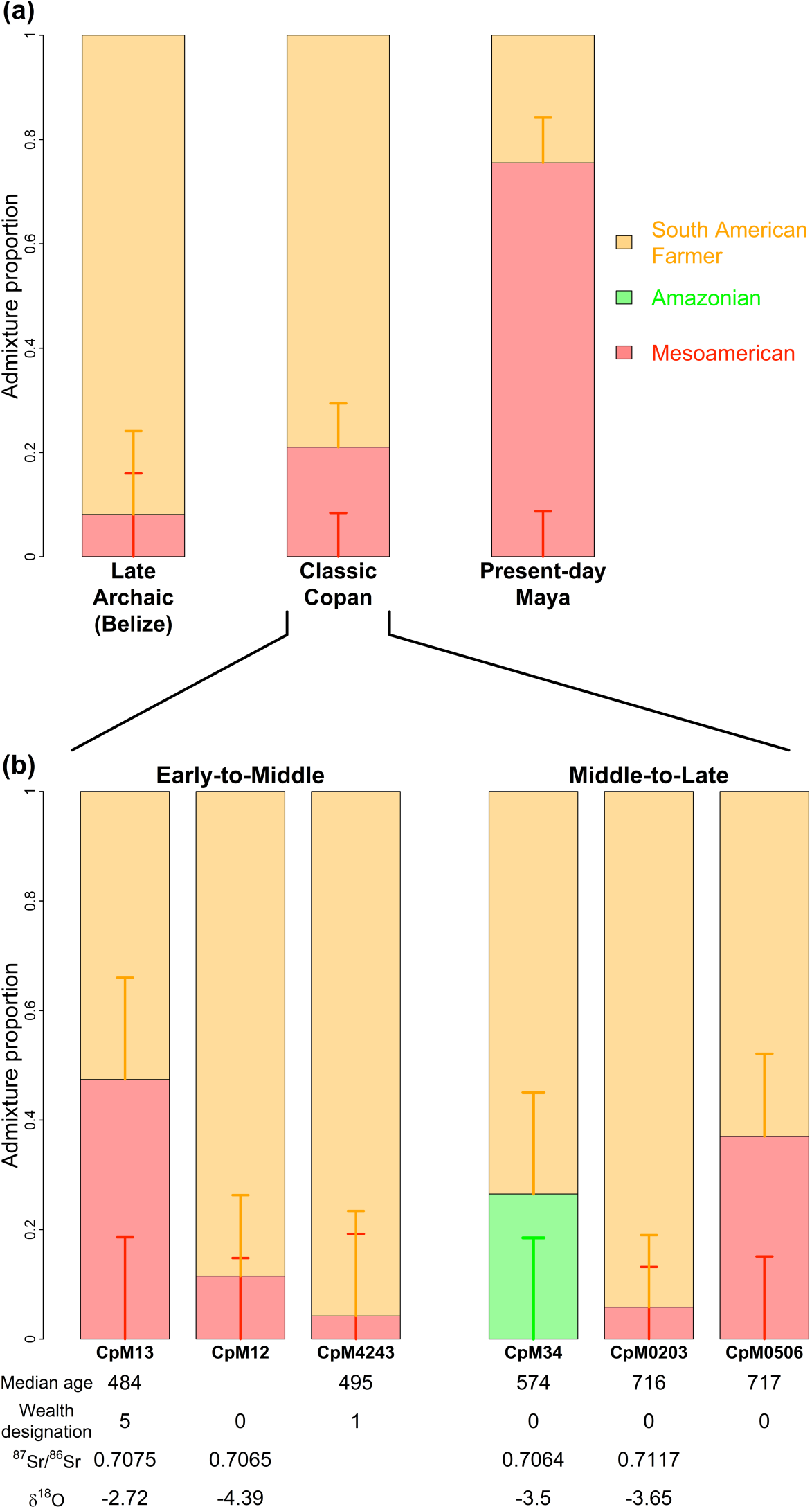
Temporal shifts in genetic ancestry (a) across the Late Archaic, Classic Copan, and contemporary periods and (b) within the Classic Copan period. Bar plots show proportions of distinct genetic ancestors: South American Farmer (orange; represented by Aconcagua and COR or LUK), Amazonian (green; represented by Surui), and Mesoamerican (red; represented by Mixe). Vertical bars represent ±1 SE estimated by *qpAdm*. The values of admixture proportions, as well as fittings of different models, are shown in tables S6 and S8. Additional information on the Classic Copan individuals are shown at the bottom if it is available, including median ages, wealth designation, and isotopic profiles.

We then investigated a potential genetic link between the Classic Copan and a group of ancient individuals who are associated with the Late Archaic in Belize (*25*). Our admixture analysis with a single ancestry model confirms that the genetic makeup of Classic Copan can be sufficiently explained by Late Archaic ancestry (*p* = 0.179), indicating a certain level of genetic continuity. This genetic link between the Copan and Late Archaic is also observable as a shared ancestral component in the ADMIXTURE profiles (fig. S6). We find that the two-way admixture of Mesoamerican and SAF ancestry fits the Late Archaic population (*p* = 0.117). However, the admixture proportions are highly skewed towards SAF ancestry (>90%), with a large standard error of ± 16.0% (Figure 4a and table S6). Therefore, the Mesoamerican component in the LateArchaic can be considered extremely low, which is also supported by a nested model selection, where SAF adequately represents genetic ancestry of the LateArchaic (*p* = 0.589; see table S6). These results suggest the Late Archaic period marks the emergence of a dual ancestral structure in the region.

### Genetic continuum with dual ancestry across the Classic Copan individuals

The two-way admixture of Mesoamerican and SAF ancestry fits five Copan individuals (tail probability > 0.05), who ranked the Mayan as the top populations in the *f*_4_-tests (fig. S7), illustrating varying levels of admixture proportions (Figure 4b and table S8). CpM0506 and CpM13 demonstrate intermediate proportions of the two distinct ancestors, with 37.0% ± 15.1% and 47.4% ± 18.6% Mesoamerican ancestry, respectively (table S8). We find that the admixture model fits those two individuals significantly better than either single-ancestry models (*i*.*e*., Mesoamerican or SAF only) (*p*-values for the nested models < 0.01). These Mesoamerican components in CpM0506 and CpM13 are also observable in their ADMIXTURE profiles (fig. S6). In contrast, the remaining individuals, CpM0203, CpM12, and CpM4243, exhibit nearly or more than 90% of SAF ancestry. Due to these skewed ancestry proportions and possibly also the limited genomic data coverage, SAF ancestry alone can sufficiently explain the genetic makeup of these three individuals (nested *p*-values > 0.05; table S8).

Different from those Classic Copan individuals, CpM34 shows a strong genetic affinity to LUK, one of the ancient Andean populations (fig. S7). Our PCA and ADMIXTURE results confirm that LUK are genetically clustered with the other ancient Andean populations (Figure 1; figs. S4 and S6), all of which share SAF ancestry. This finding is further supported from our *f*_4_-tests with the form of *f*_4_(Mbuti, *X*; *Y*, LUK), where Rio_Uncallane, who are 1,800-years-old sedentary farmers from Bolivia (*28*) and uniquely characterised by SAF ancestry (Figure 2; fig. S6), emerges as the top population exhibiting genetic affinity to LUK (fig. S11). Applying the *f*_4_-tests to Chane and Piapoco, the second and third top-ranked populations in our *f*_4_(Mbuti, *X*; *Y*, CpM34) (table S2), we find both populations have strong genetic links with Surui, a present-day Amazonian population from Brazil. We then modelled CpM34 by a two-way admixture between LUK, representing SAF ancestry, and Amazonian ancestry (table S8). Although the admixture model fits this individual, ∼70% of genetic ancestry derives from SAF, making the SAF-only model the most parsimonious (nested *p*-value = 0.063). Overall, these results highlight a potential genetic cline of Mesoamerican ancestry at the individual level.

## Discussion

Our newly generated genomic data of seven individuals associated with the Classic Maya culture provide compelling evidence of a dual-ancestral structure of the Copan kingdom, characterized by both Mesoamerican and SAF. Our results from dating and modelling of the admixture events support an idea that this dual structure may have emerged during the Late Archaic period (*25*). This period saw the intensified reliance of maize in Central America (*32*), which could have been reintroduced from South America after maize dispersed into the south and then became fully domesticated (*33*, *34*). This transformative process, marked by south-to-north population movements, potentially explains the prevalence of SAF in the region during the Late Archaic period. In the Classic period, Mesoamerican ancestry became more pronounced, in contrast to the Late Archaic period, resulting in a significant increase of this ancestral legacy from the Classic period onward.

Our analysis clearly demonstrates varying proportions of distinct genetic ancestry among the individuals, which can be linked with archaeological and isotopic contexts. The Copan individuals are chronologically divided into two groups (Table 1 and Figure 4): the Early-to-Middle Classic and the Middle-to-Late Classic. The enigmatic dynastic ruler, CpM13, exhibits nearly equal proportions of the two ancestors, with 47.4% Mesoamerican and 52.6% SAF ancestry. The strontium and oxygen isotope ratios from this individual (Table 1) strongly suggest an origin as a likely immigrant from a lowland area, rather than being locally rooted in the highland Copan region (*14*). The sacrificial person of CpM12, discovered in close proximity to the dynastic ruler, is genetically identified as SAF. The isotopic profile of this individual supports a local origin (*14*), implying that SAF ancestry could have been already present in the Copan state. This is further corroborated by another individual within the same chronological group, CpM4243, whose genetic ancestry also traces back to SAF origins.

Two out of three individuals from the Middle-to-Late Classic group, CpM34 and CpM0203, trace their genetic roots to SAF ancestry (Figure 4). Both individuals are archaeologically classified as low rank. Their oxygen isotope ratios support an origin from highland areas, with CpM34 potentially being local, as indicated by the strontium isotope ratio falling within the baseline range in Copan (0.7052 - 0.7072) (*12*). However, CpM0203 exhibits an unusually high strontium isotope value (0.7117); such elevated values are observable in the Maya Mountains of southern Belize (*14*), the Guatemala Highlands (*35*), or western Honduras (*36*) within Central America. Alternatively, this strontium isotope ratio may indicate an origin in highland areas in South America (*37*), consistent with its own genetic ancestry of SAF. However, due to a lack of baseline measures in other parts of Central America, including the region spanning from El Salvador to Colombia, where archaeological, anthropological, and linguistic ties with Classic Copan have been proposed (*7*), the geographic origin of this individual remains unclear. Our analysis identifies another individual with dual-ancestry from the Late Classic group, CpM0506, although archaeological and isotopic details are unavailable for this individual. Despite the variation observed in the genetic and geographic origins of the Classic Copan individuals, all of them relied on maize as a primary dietary source (fig. S13), aligning with findings from previous studies (*32*, *38*).

An in-depth profiling of funerary contexts associated with Maya culture reveals nearly 90% are classified as low-rank (*i*.*e*., scores of “0” or “1”), with only 5% assigned to scores of “4” or “5” (*i*.*e*., high-lank) (*39*), suggesting that the majority of the populations fall within the low-rank category. Within our sampled individuals ranked either “0” or “1” (Table 1), all show their genetic origins in SAF (Figure 4). Notably, two individuals among them are likely to have originated locally. In contrast, the dynastic ruler, who, like the first king (K’inich Yax K’uk’ Mo’) (*40*), was born and grew up elsewhere before relocating to Copan, exhibits a distinct composition, featuring Mesoamerican as well as SAF ancestry. Given the potential prevalence of SAF ancestry before the Classic period (Figure 4), the Mesoamerican components in the ruler may signify the legacy of royal families.

There are caveats to this analysis. Firstly, our study is constrained by the small sample size, involving only seven Classic Copan individuals, the majority of whose genomes have low coverage. These are well-known limiting factors in statistical tests and admixture modelling (*41*). Secondly, our sampling is non-random, focusing on individuals from the same residential or patio groups (Table 1). Considering the observed archaeological and isotopic variation across Copan sites (*12–14*), a more extensive and denser sampling of ancient genomes will be essential to trace the temporal and regional diversity of the population within the state, which will then provide a more comprehensive understanding of the dual-ancestral structure proposed here for the Classic Copan.

In summary, our research offers a new perspective, suggesting the existence of broader mobility and social networks beyond the conventional boundaries of the Maya territory and the homelands for migrants within Central America. This underscores the significance of the Copan kingdom as a crossroads, facilitating connections between Central and South America. Ancient genomics on the Classic Copan provides a unique opportunity to integrate a genetic ancestry continuum into the widely accepted archaeological concept of the dichotomy between local and non-local or between Maya and non-Maya—commonly acknowledged in the understanding of the origins of people in the Maya states.

## Materials and Methods

### Sampling

Petrous bones from a total 16 samples (*i*.*e*., petrous bone fragments) excavated from the sites of Copan in western Honduras were sampled and brought for processing at Kanazawa University in Japan. Detailed information on the archaeological contexts is provided in note S1.

### DNA extraction

Sample processing was carried out in a dedicated ancient DNA facility at Kanazawa University where all the precautions for ancient DNA processing were followed as described in a previous study (*42*). Samples were photographed extensively before further processing. All bones were exposed to ultraviolet light for 15 min on either side to remove surface contaminants. Further cleaning of their surface was performed with a drill before extraction. The otic capsule region of the petrous temporal bone was targeted for sampling. An aliquot of 40-100 mg of the bone powder was subjected to a silica column method with an initial washing step by predigestion solution (*43*). DNA extracts were purified with MinElute silica columns (QIAGEN) and eluted at a volume of 30 μl of the elution buffer.

### Library preparation and sequencing

The initial screening of each sample and blank controls was performed by constructing a double-stranded DNA next-generation sequencing library mainly using a NEBNext Ultra II DNA Library Prep Kit for Illumina with bead-based size selection. Every library was screened on a MiSeq Illumina platform. The samples with endogenous contents between 1% and 17% were selected for high-coverage sequencing and subsequently treated with uracil-DNA-glycosylase (UDG) before a second library construction (*44*). The concentration and quality were then assessed using the Agilent TapeStation 4200 system with the High Sensitivity D1000 ScreenTape. High-coverage sequencing for UDG-treated libraries from seven individuals was carried out on a NovaSeq 6000 Illumina platform (150-bp paired-end reads) at Macrogen (Republic of Korea).

### Sequence data processing

We trimmed adapters using *AdapterRemoval* version 2.2.2 (*45*), with the following parameters: --collapse --minadapteroverlap 1 --minlength 25 --minquality 25 --trimqualities. The trimmed reads were aligned to the hs37d5 human reference genome using *BWA* (Burrows-Wheeler Aligner) version 0.7.5 (*46*), with relaxed parameters, “bwa aln -l 16500 -n 0.01 -o 2.” We used *picard-tools* version 2.22.1 (https://broadinstitute.github.io/picard/) to add reads groups, remove PCR duplicates, and merge all different libraries sequenced on MiSeq and NovaSeq platforms into a single BAM per individual. The authenticity of ancient DNA was determined in non-UDG-treated libraries by looking at degradation patterns at 5′- and 3′-end, respectively, using mapDamage2.0 (*47*). The reads were further filtered for the length longer than 34 bp and mapping quality higher than 25. Two final processing steps were carried out on the merged files: Indels were realigned using Genome Analysis Toolkit (GATK) version 3.7-0 (*48*), and the quality of the two bases at the end of each read was manually reduced to a score of 2 (“softclipped”), to avoid calling genotype from these damage-prone regions. Genomic coverage and depth were measured using *qualimap* version 2.2.1 (*49*). We used the same processing pipeline for published ancient genomes in FASTQ format with the exception of using cutadapt version 1.15 (*50*) instead of AdapterRemoval. BAM files were realigned to our reference genome and then put through the same pipeline. Quality assessment of raw data was conducted with FastQC v0.11.4 (*51*).

### Relatedness and molecular sex determination

The kinship was estimated for ten samples that were sequenced to high-coverage using “Relationship Estimation from Ancient DNA” (READS) (*52*). This analysis identified three pairs of samples as genetically identical (CpM02-CpM03; CpM05-CpM06; CpM42-CpM43; see also note S1), which were subsequently merged into single individuals of genomes (*i*.*e*., CpM0203, CpM0506, and CpM4243). We then re-ran READS for seven Classic Copan individuals. Any first- or second-degree relatives were removed from analysis. To determine the sex of each of our ancient samples, we filtered for reads with a minimum mapping quality of 30 and followed the method outlined in a previous study (*53*). In summary, we calculated the ratio of Y chromosome reads to the total number of sex chromosomes (*R_y_*) and assigned female if *R_y_* < 0.016 or male if *R_y_* > 0.075.

### Contamination and mitochondrial DNA haplogroups

To assign mitochondrial (mt) haplogroups, we extracted sequence reads uniquely mapped to mtDNA from BAM files using *SAMtools* (version 1.7) (*54*) and re-aligned the reads to the revised Cambridge reference sequence using *BWA* (version 7.5a) (*46*). Consensus sequences were determined to mapping quality higher than 25, a minimum depth of consensus of 5, and a base quality of 30 using mpileup and bcftools within the SAMtools package. We then assigned a specific haplotype to each consensus sequence using *HaploGrep* (version 2.2.9) (*55*).

We estimated contamination rates as previously described in (*56*). Briefly, we calculated the rate of secondary bases that did not match the consensus sequence at haplotype-defining and private mutations defined by *HaploGrep*.

### Y-chromosome haplogroup determination

All newly sequenced individuals determined to be male were piled up to the 2019 version of the International Society of Genetic Genealogy database using GATK version 3.7-0 (*48*), with a mapping quality of 20 and a base quality of 30.

### Genotype calling and merging ancient with present-day genomes

We created a large panel that includes newly sequenced Classic Copan genomes and publicly available ancient genomes from Siberia and the Americas (see table S1 and fig. S3). We genotyped all ancient individuals for the biallelic SNP sites present on two different reference panels, to which they were subsequently merged: (i) the Human Origin Array (HOA) consisting of 594,896 SNP sites for 1,963 modern, ancient, and reference genomes (*57*) and (ii) the Simon Genome Diversity Project (SGDP) consisting of 278 modern, ancient, and reference genomes (*58*) that have been filtered for autosomal transversion-only SNPs with a minor allele frequency of 1%, which left 3,867,550 SNP sites in our merged data. For each SNP site, we randomly called a high-quality single base (bq30) per position to create a pseudo-diploid genotype using GATK version 3.7-0 (*48*). All populations from the Americas on the HOA dataset or Mayan on the SGDP were masked for genomic regions that were identified as inherited from non-American populations (*i*.*e*., those from Africa or WestEurasia) by MOSAIC (*59*), with the following parameters “--ancestries 3 --panels Africa America Siberia EastAsia SouthAsia WestEurasia.” Genomic coordinates of non-American segments were obtained with the R function “grid_to_pos()” included in the MOSAIC package, converting grid points based on recombination distances into physical positions.

### Principal Component Analysis

PCA was conducted using smartpca (v16000) from the EIGENSOFT package (v7.2.0) (*60*). All Classic Copan and a subset of other ancient individuals were projected onto present-day Americans in the HOA dataset. The options specified in our runs are as follows: “killr2: NO,” “numoutlieriter: 0,” “lsqproject: YES,” “shrinkmode: YES.” We excluded ancient individuals if the number of missing sites is larger than 10,000.

### ADMIXTURE

Unsupervised genetic clustering was conducted by ADMIXTURE version 1.23 (*61*), with the same set of ancient and present-day individuals as used in PCA. SNPs were ascertained for sites genotyped at least in one individual of the Classic Copan, leaving 386,696 SNPs for analysis. We ran 100 iterations with random seeds for the number of clusters (*K*) from 2 to 7 and chose the run with the minimum cross-validation error for plotting.

### F4-statistics

We used qpDstat (version 900) for the merged datasets of ancient individuals and the SGDP panel of present-day populations, with the *f_4_* mode in the AdmixTools v6 package (*62*). Mbuti was always specified as an outgroup, and standard errors in *f*_4_-statistics were calculated from the default block jackknife approach. Statistical significance in genetic affinity between populations was defined as *Z*-score greater than 2.5 or 3.0 if the test was applied to a specific individual or a group of individuals.

### Admixture modelling by qpAdm

To model admixture events in the Classic Copan and other ancient and present-day populations from the Americas, we applied qpAdm v1000 in the AdmixTools v6 package (*62*, *63*) to the merged data of ancient and SGDP populations. Our analysis only used transversion sites with global minor allele frequencies of >1%, coupled with the option of “allsnps: YES.” A set of 13 ancient and present-day populations was used as an outgroup in the modelling: Han (*n* = 3), Pima (*n* = 2), Karitiana (*n* = 3) (*58*), MA1 (*n* = 1) (*64*), USR1 (*n* = 1) (*65*), ASO_4000 (*n* = 1) (*66*), Sumidouro5 (*n* = 1), Ayayema (*n* = 1), PuntaSanataAna (*n* = 1), SpiritCave (*n* = 1) (*29*), Anzick (*n* = 1) (*67*), and Taino (*n* = 1) (*68*). We considered that an admixture model fits data if a tail probability is greater than 0.05.

### Dating admixture events by DATES

We used DATES v753 (*31*, *69*) to estimate the time of admixture between Mesoamerican and South American Farmer ancestry in the Classic Copan. Mixe was used to represent Mesoamerican ancestry, while Aconcagua and COR were used as the second source. The estimated date in generation was converted into years, assuming a generation time of 25 years, which was further added into a mean age of mediate dates from six Classic Copan individuals (*i*.*e*., 1,353 years before present). The parameter settings that we used are as follows: binsize: 0.001, maxdis: 1.0, runmode: 1, mincount: 1, and lovalfit: 0.45. The SE was estimated from a weighted block jackknife method.

## Acknowledgments

We would like to thank Honduran Institute of Anthropology and History (IHAH), particularly Dr. Rolando Canizales Vigil (General Manager), Omar Talavera (Sub-Manager of Patrimony), Salvador Varela (Regional Director of Western Region), and Carlos Carbajal (Administrative Co-Director of PROARCO). We also acknowledge the support of Komatsu University, specifically Dr. Hiroshi Yamamoto (President) as well as the administrative support staff, including Yoichi Sato and Miguel Echeverria.

## Author contributions

S.Nakamura, T.G., and S.Nakagome supervised the study. S.Nakamura, M.F., and M.O. excavated the sites, provided samples, and assembled archaeological and anthropological information. T.G. performed ancient DNA laboratory analysis. M.M. and S.Nakagome processed and analysed genetic data. M.M., S.Nakamura, T.G., and S.Nakagome wrote the manuscript with input from all co-authors.

## Data and materials availability

Genome sequence data produced in this study are available at the European Nucleotide Archive (ENA) with accession number XXX. All data needed to evaluate the conclusions in the paper are present in the paper and/or the Supplementary Materials.

