## Supplementary Materials for "Ancient genomics reveals a genetic continuum with dual structure in the Classic Copan"

### Supplementary Text

#### Note S1. Archeological contexts of Copan

The ancient city lies in the Copan valley, in western Honduras, at an altitude of approximately 600 metres (*70*). By 1,000 BCE, the villagers of Copan already practised agriculture and constructed elevated stone platforms for their living quarters and for burying their dead underneath it. By 300-400 CE, the small settlement had become a vibrant city with economic opportunities for a growing population (*7*). In 427 CE, arrived *K'inich Yax K'uk' Mo'*, the dynastic founder of Copan, who later commissioned the first monument of the valley in the Classic Maya style with according hieroglyphic inscriptions (*70*). He erected the original buildings of the Acropolis, which his successors rebuilt and extended over time into a monumental complex (*7*). The Acropolis was the heart of political, economic, and religious life in Mayan civilization (*71*), encapsulating the East and the West courts, the three hectare Acropolis of Copan accommodates temples, administrative buildings, and the royal residence. Like most buildings in the Acropolis, the ball court filled in both religious and political roles as the outcome of the game could decide on religious actions to take, and even the outcome of a battle. Building facades are decorated with high-relief sculptures in the 8th century, and many stelae and altars ornate the plazas (*7*, *71*). At 757 CE, the 15th ruler finished one of the most impressive monuments of Copan, the Hieroglyphic Stairway, which represents today the longest pre-Columbian inscription in the Americas (*70*).

***Nomenclature***

To date, more than 3,500 structures have been confirmed in the Copán Valley, where a total area of approximately 24 $km^2$ is organised into 500-metre grid squares in order to identify and register each of these structures. Each square block is designated by a combination of a number and a letter, indicating its vertical and horizontal axes respectively (*e.g.*, 10L, 9M, or 8N). Within a square block, different structures are numbered sequentially, starting from one. For example, a structure labelled as "10J-45" represents that it is the 45-th structure within the block marked as 10J. This nomenclature  makes it possible to locate any structure on the archaeological map of the Copán Valley, published in 1979 (*72*).

***Archaeological contexts of seven Classic Copán individual sequenced in this study***

The structures, from which the samples sequenced for high-coverage were excavated, are registered as two distinct residential clusters. The first cluster lies approximately one kilometre west of the Acropolis, specifically within the 10J block, comprising Structures 10J-39, 10J-45, and 10J-71. The second cluster, located north of the Great Plaza, falls within the 9L block and consists of structures 9L-104 and 9L-105 (see maps below).

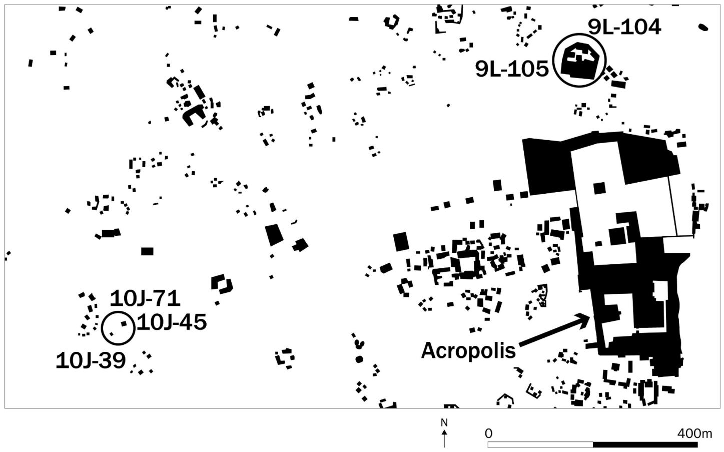


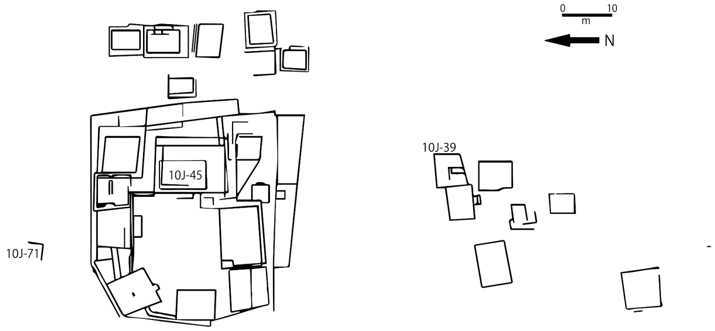


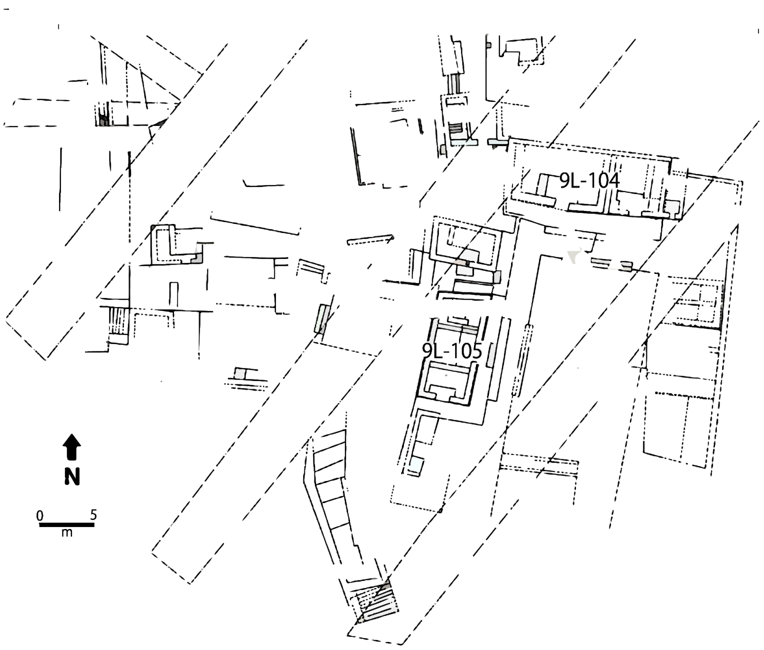


It is important to note that there is a discrepancy between the wealth designation scores assigned to individuals and their archaeological contexts. These two structural clusters are located in close proximity to the core of Copán, namely the Great Plaza and Acropolis, often indicating their significance in the Classic Copán state. Therefore, the inhabitants of these structures can be regarded as part of the elite. However, it is worth mentioning that the wealth designation measures are based on the criteria outlined by Krejci and Culbert (1995) (*73*), which may not be fully applicable to burial contexts in the ancient Copán society.

***CpM0203 and Cp0506***

Burial 7-2000, located within the interior of the southeast corner of Structure 10J-39, was discovered in 2000, as part of the Integral Program for the Conservation of Copan Archaeological Park (Programa Integral de Conservación del Parque Arqueológico Copán, or its acronym in Spanish, PICPAC). This archaeological project was conducted by the Honduran Institute of Anthropology and History (IHAH), under the direction of S. Nakamura with the assistance of M.F. There were human remains belonging to two distinct individuals; neither individuals were accompanied by any artefacts as offerings. Burial 7(1) was identified as a primary *in situ* burial during field work, whereas Burial 7(2) was subsequently recognized in a laboratory setting due to the presence of identical bone elements. The remains of Burial 7(1), designated as CpM0203, were well-preserved, comprising an almost complete adult skeleton. In contrast, the remains of Burial 7(2) (CpM0506) were fragmented. As evident in the photograph taken by S. Nakamura (see a picture below), only a portion of the skull from CpM0506 was found resting beside the waist of CpM0203. This burial arrangement suggests that CpM0506 was a secondary burial accompanying CpM0203. While this observation led us to hypothesise a potential kinship between these two individuals, our analysis demonstrates that there is no familial relationship between them, at least up to the second-degree level.

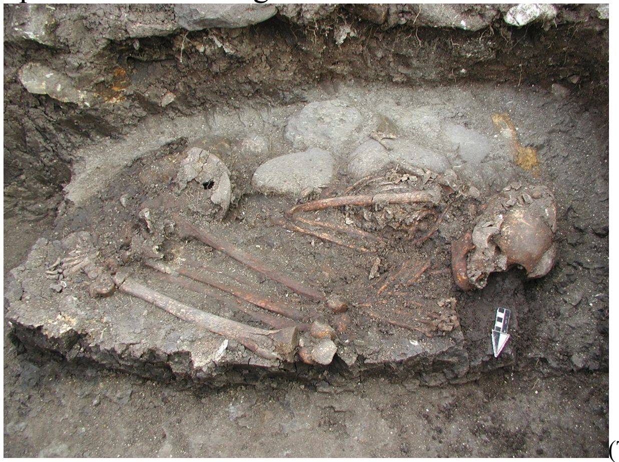

(The photo was taken by S. Nakamura)

***CpM11***

Burial 27-2000 was uncovered during the PICPAC excavation in 2000. The remains of an individual, CpM11, were discovered *in situ* outside the southern wall of Structure 10J-71, at the base brick-line level. While the remains are generally well-preserved, they were fragmented but identified as an adult. Due to the presence of a jade bead left as an offering, this individual was categorised as level 1 in terms of wealth designation.

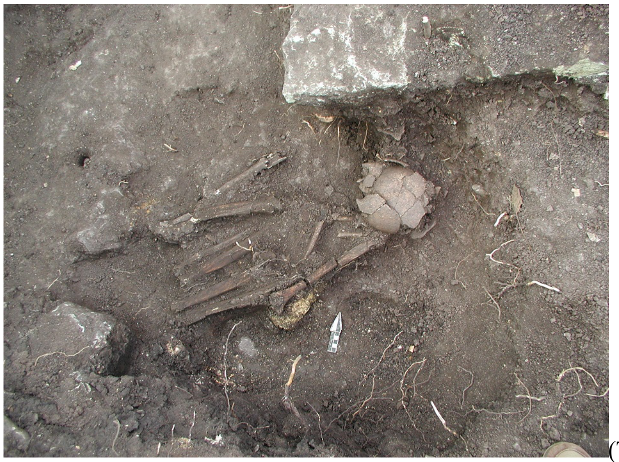
(The photo was taken by S. Nakamura)

***CpM12***

Burial 35-2000 was excavated as part of the PICPAC in 2000, exhibiting the well-preserved remains of an adult. The human remains, designated as CpM12, were discovered *in situ* directly beneath the first brick-line of the west-facing wall (front) of the foundation of 10J-45-2nd, approximately 20 to 40 cm below the surface, without any formal delineation for burial. Structure 10J-45-2nd was constructed as an adoration shrine following the presumed burial of a governor (CpM13 from Burial 36-2000). The positioning of CpM12, with the right arm placed in front of the chest and legs crossed as if seated (see top picture below), in close proximity to the burial chamber of Burial 36-2000, suggests that this individual was the sacrificed companion of the presumed ruler, CpM13. Additionally, one of the jade pendants found within the offering box (5-2001), unearthed near Burial 35-2000, depicts a figure displaying a gesture of respect and subordination towards another individual, potentially the ruler of Burial 36-2000 (CpM13) (see bottom picture below). This gesture is reminiscent of the burial posture observed in CpM12. Contrary to early studies that concluded CpM12 was female based on morphological characters, our genomic analysis clearly identifies this individual as male. This finding underscores the importance of re-evaluating the determination of sex for royal individuals excavated from the Acropolis using genomic techniques.

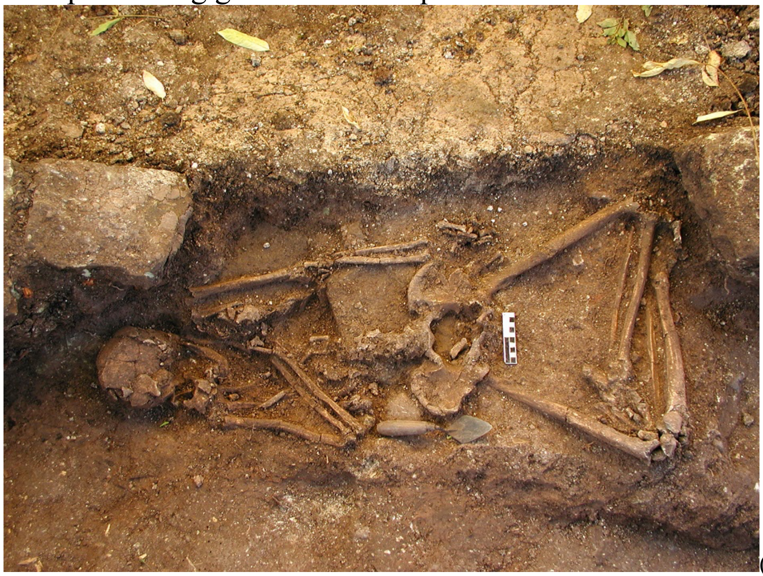
(The photo was taken by S. Nakamura)

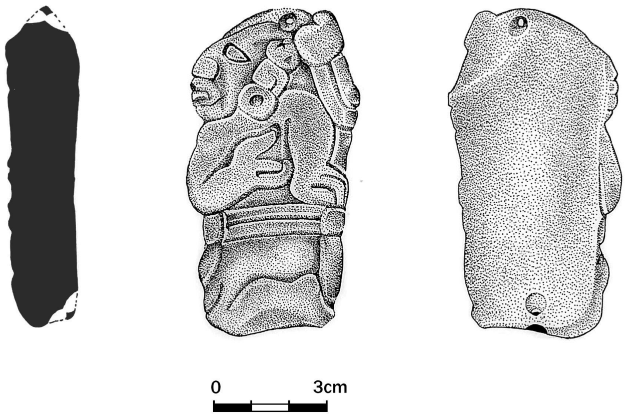

(The picture was made by PICPAC)

***CpM13***

Burial 36-2000 represents one of the largest and most significant burials uncovered outside the Acropolis in Copán's 130-year history of archaeological investigation. The excavation, conducted as part of the PICPAC project, was led by S. Nakamura and M.F. in September 2000. The exceptional aspect of Burial 36-2000 lies in its contradiction of a commonly accepted hypothesis that Copan rulers were exclusively buried within the Acropolis, the political, economic, and ritual hub of ancient Copan society.

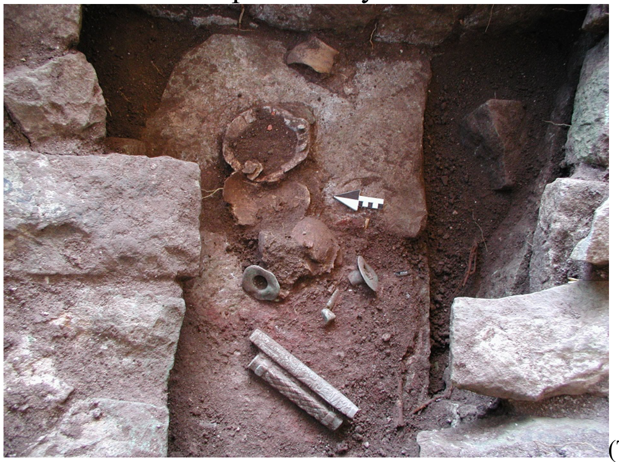

(The photo was taken by S. Nakamura)

The body remains of CpM13 are in varying states of preservation, ranging from regular to poor; they consist of fragmented and incomplete remains of an adult. Initially presumed to be a secondary burial, further analysis uncovered this individual to be the primary burial *in situ*. Red pigments were discovered on the surface of the bones. The tomb exhibited the characteristic vault of a royal tomb and contained two enormous jade pectorals (see pictures below), one measuring 20 cm in length and the other 24 cm, adorned with engraved designs symbolising the political and military authority of the individual. The archaeological evidence strongly suggests that this individual was one of the dynastic rulers of the fifth century.

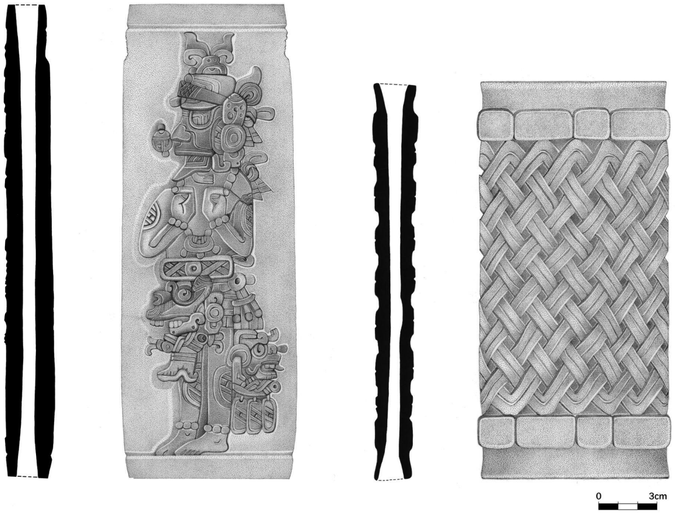


(The picture was made by PICPAC)

***CpM34 and CpM4243***

Burial 94 and Burial 157, located in Structure 9L-104 and Structure 9L-105 respectively, were excavated in 2007 and 2016 as part of the new Copan Archaeological Project (known by its acronym in Spanish: PROARCO, Proyecto Arqueológico Copán). PROARCO was conducted by the Honduran Institute of Anthropology and History (IHAH), under the direction of S. Nakamura with the assistance of M.F. These two structures were found in group 9L-23, also known as the Núñez-Chinchilla group, located 150 metres north of the Great Plaza.

Burial 94 contained the fragmented and incomplete remains of a girl (CpM34; left picture), in a poorly preserved state. Archaeologically, it suggests that she may have been a sacrificed companion of another burial found in a box within the same structure, 9L-104. In contrast, Burial 157 consisted of the fragmented but well-conserved cranium of CpM4243 (right picture), accompanied by a ceramic offering. However, there is a possibility that this offering might have belonged to a nearby burial registered as Burial 159, one of the burial groups found in Structure 9L-105. Therefore, CpM4243 may have been one of the companions of the main burial within Structure 9L-105. According to established ceramic chronology and C14 dating results, CpM4243 is contemporaneous with Burial 36-2000, despite the considerable distance between the two structures, approximately 1 km apart.

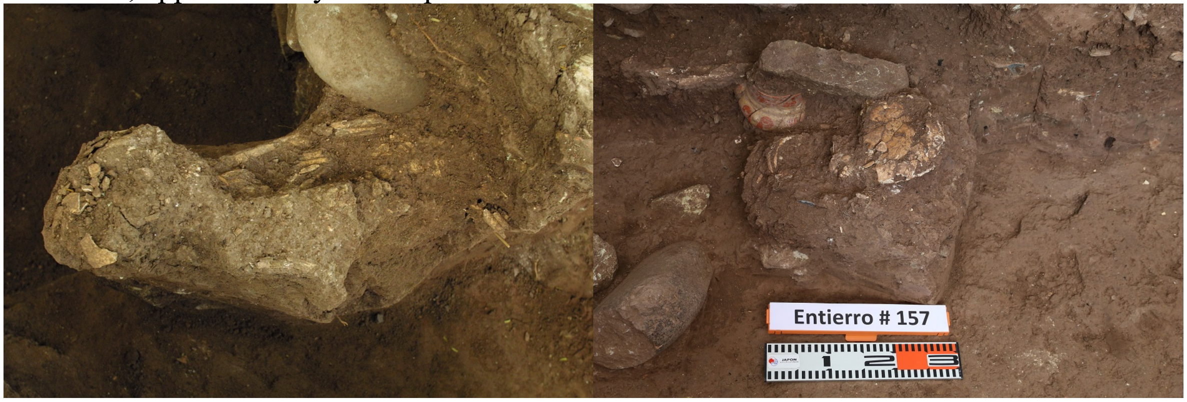


(The photos were taken by S. Nakamura)

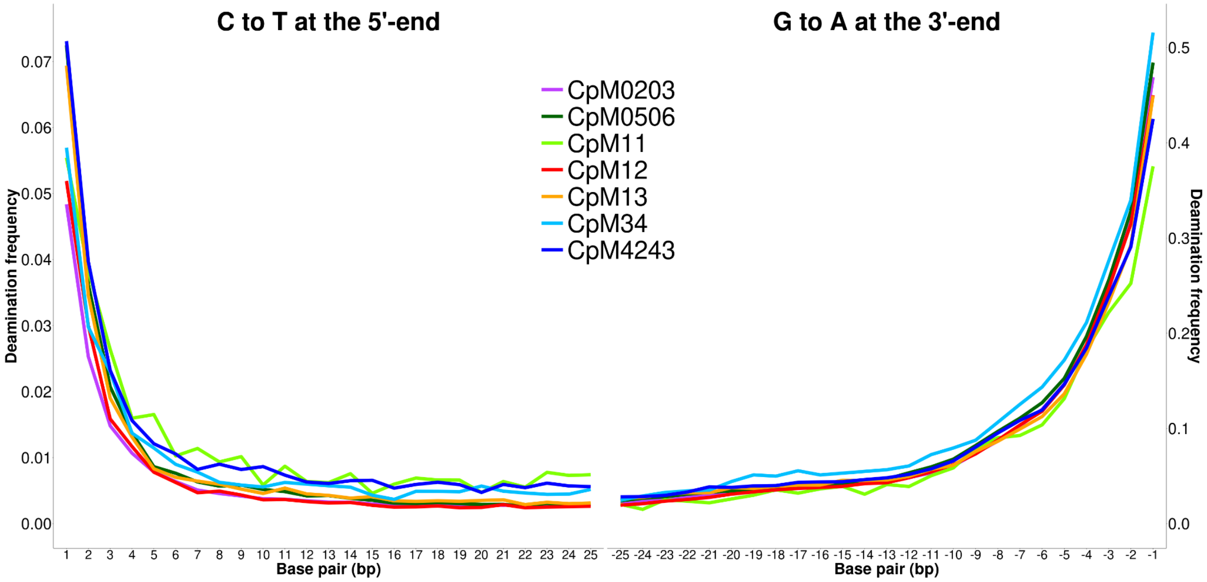


**Fig. S1. DNA damage plots for newly sequenced Classic Copan genomes.** Damage patterns on the left plot show C>T misincorporations at the 5'-end, while those on the right plots show G>A misincorporations at the 3'-end. As sequencing libraries were constructed with the NEBNext Ultra II DNA Library Prep Kit that uses USER enzyme for exercising adaptors taking a hairpin loop structure, C-to-T substitutions observable at the 5'-end become less pronounced compared with G-to-A substitutions at the 3'-end.

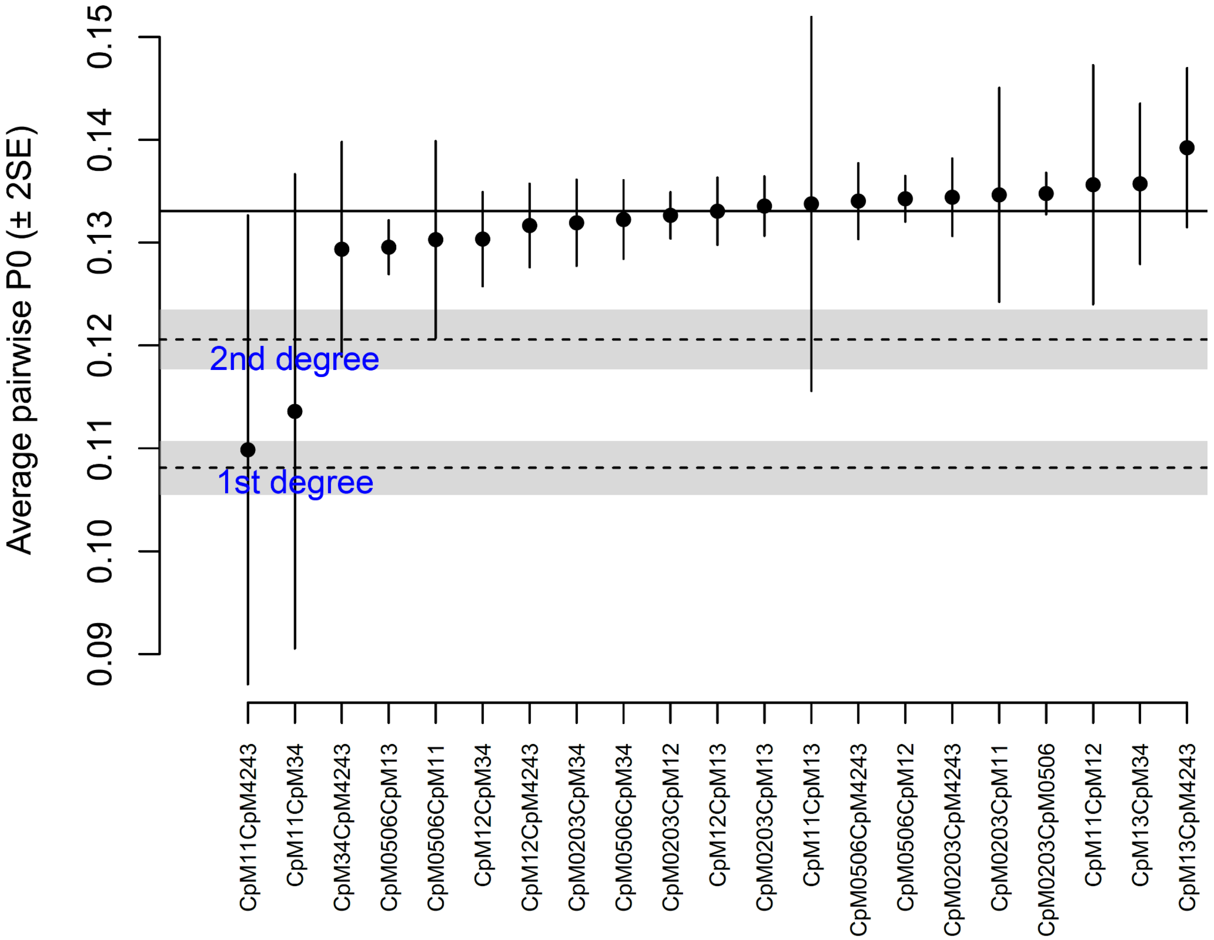


**Fig. S2. Results of kinship analysis by READ for all Classic Copan individuals sequenced in this study.** Pairs of individuals tested are listed on the *x*-axis, while the *y*-axis represents average pairwise P0 scores. Error bars show the two standard errors of the mean, and dashed lines show the cut-off for the 1st and 2nd degree relatives.

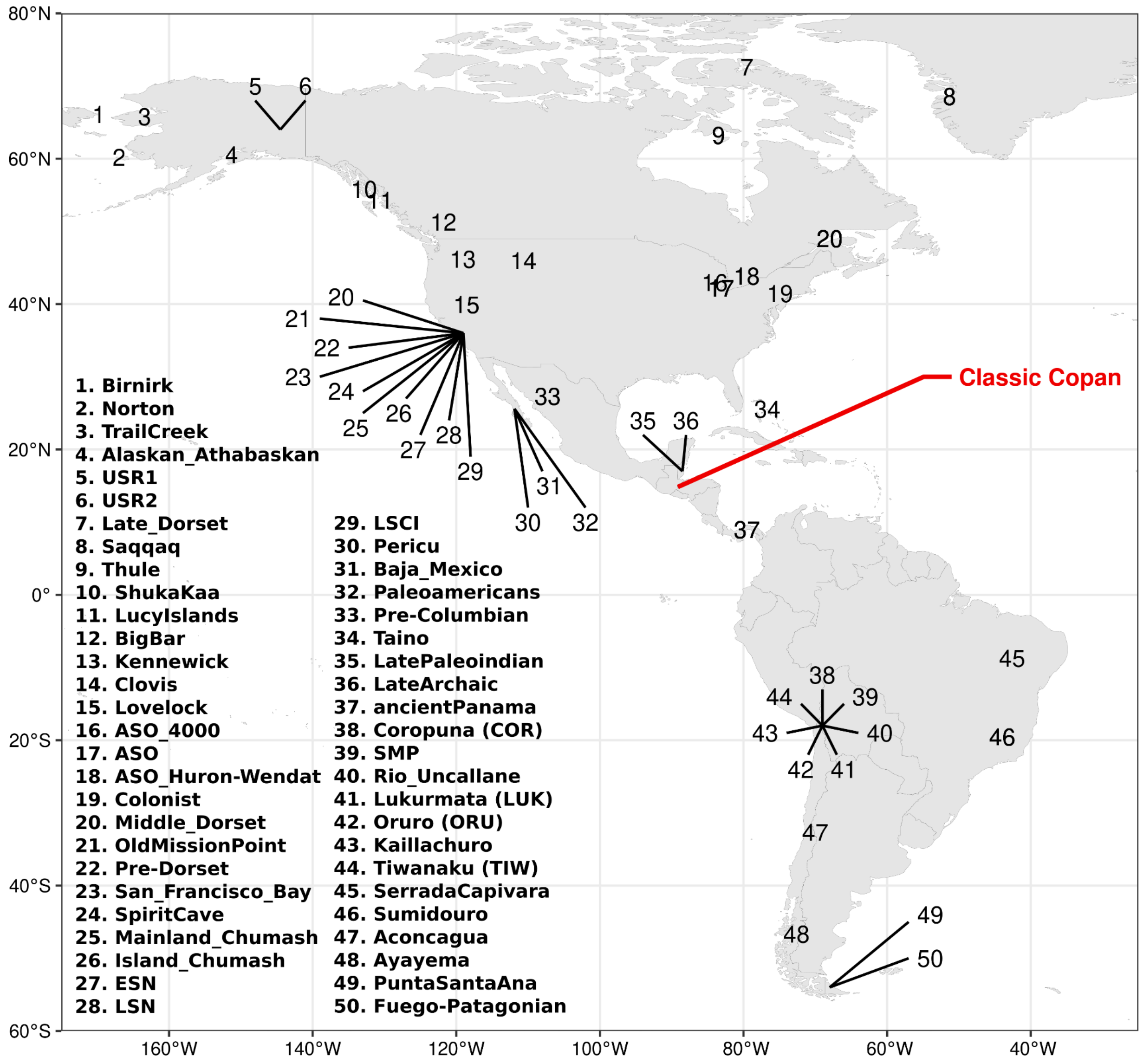


**Fig. S3. A map showing geographic locations of the Classic Copan and publicly available ancient genomes analysed in this study.** Two ancient individuals from Siberia (AG2 and MA1) included in our analysis are not plotted on the map.

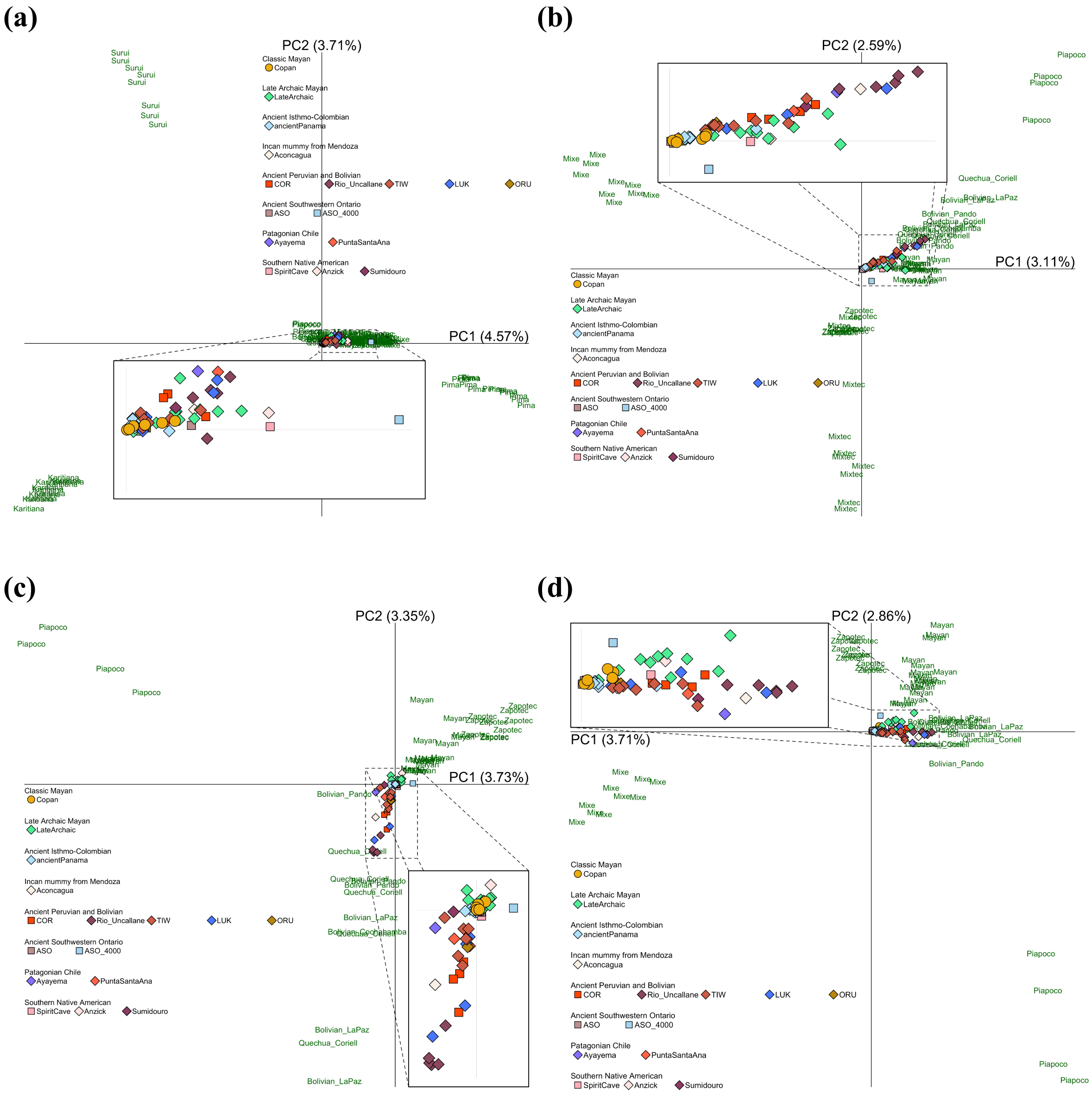


**Fig. S4**. **Principal component analysis with different sets of present-day American populations**. Ancient populations are projected onto different sets of present-day American populations: **(a)** all populations from the Americas included in the Human Origin Array, **(b)** Karitiana, Surui, and Pima are excluded from the set of populations used in (a), **(c)** Mixe is excluded from (b), and **(d)** Mixtec is excluded from (b).

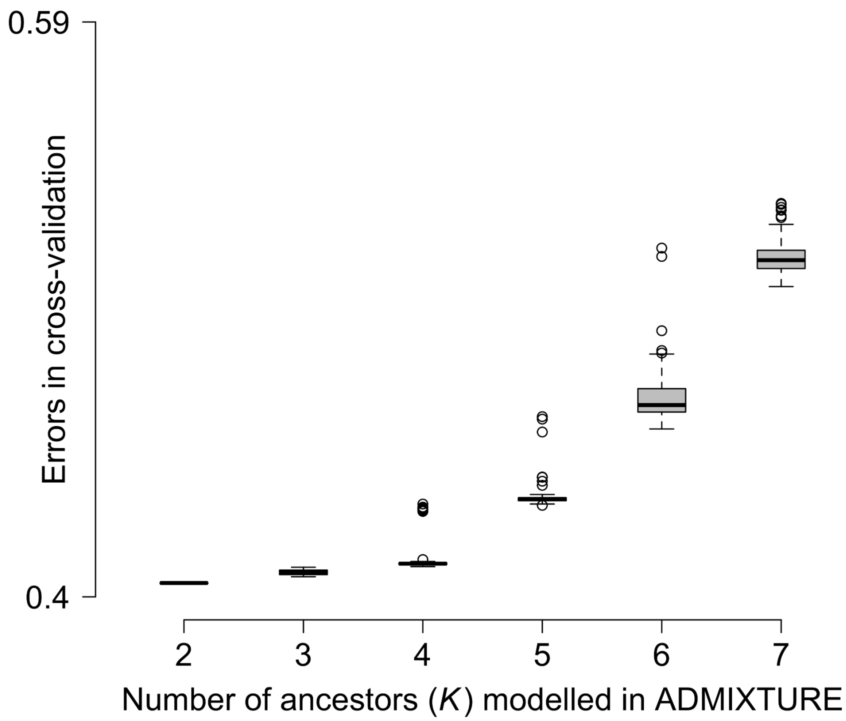


**Fig. S5. Cross validation (CV) errors used in ADMIXTURE analysis.** Each *K* component includes 100 replicates. The run with the lowest CV is plotted in Figure. 1 and fig. S6.

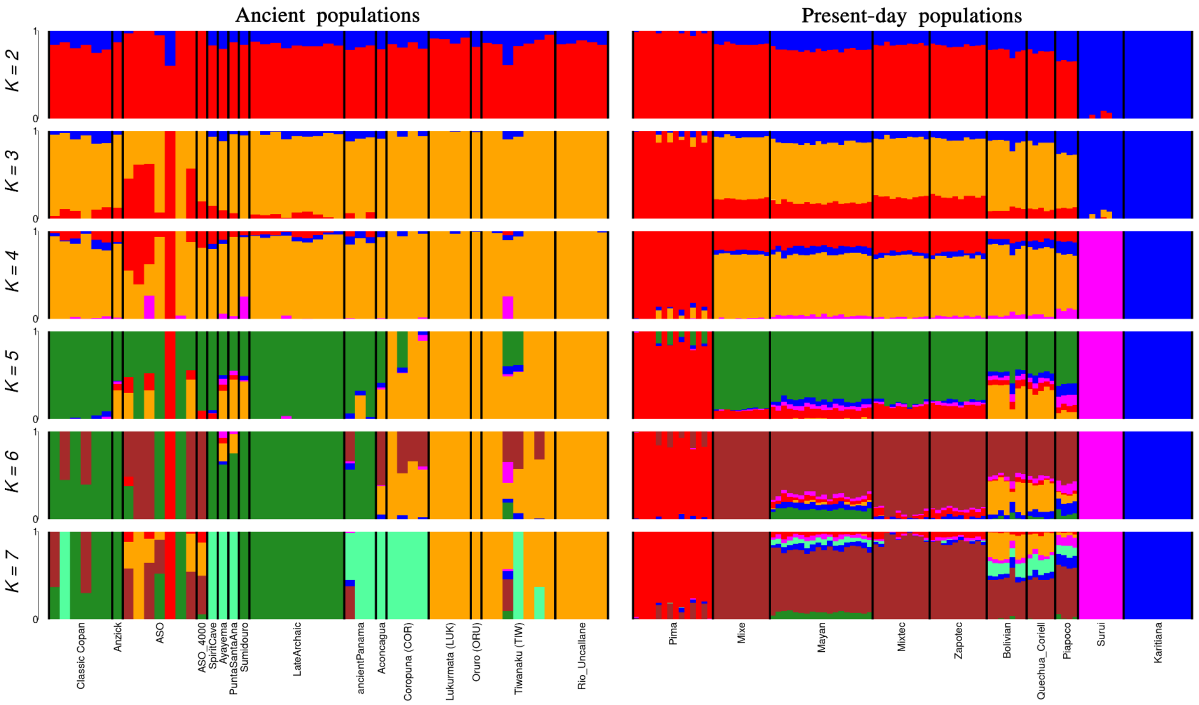


**Fig. S6. ADMIXTURE results from ancestral clusters *K* = 2 to 7 in ancient (left) and present-day (right) populations from the Americas.** Bar plots for each ancestral cluster represent the run with the lowest cross-validation error out of 100 iterations. At *K*=2, all populations separate into Amazonian and (blue) and non-Amazonian (red) ancestry. At *K*=3, the orange component emerges, which is prevalent across ancient and present-day populations but defines genetic ancestry of South American farmers (Rio_Uncallane). At *K*=4, Amazonian ancestry is divided into Karitiana (blue) and Surui (magenta). At *K*=5, the dark green component emerges, associated with LateArchaic. At *K*=6, the dataset defines Mixe (brown), and lastly, at *K*=7, Coropuna (COR) and Aconcagua are separated by the sea green component.

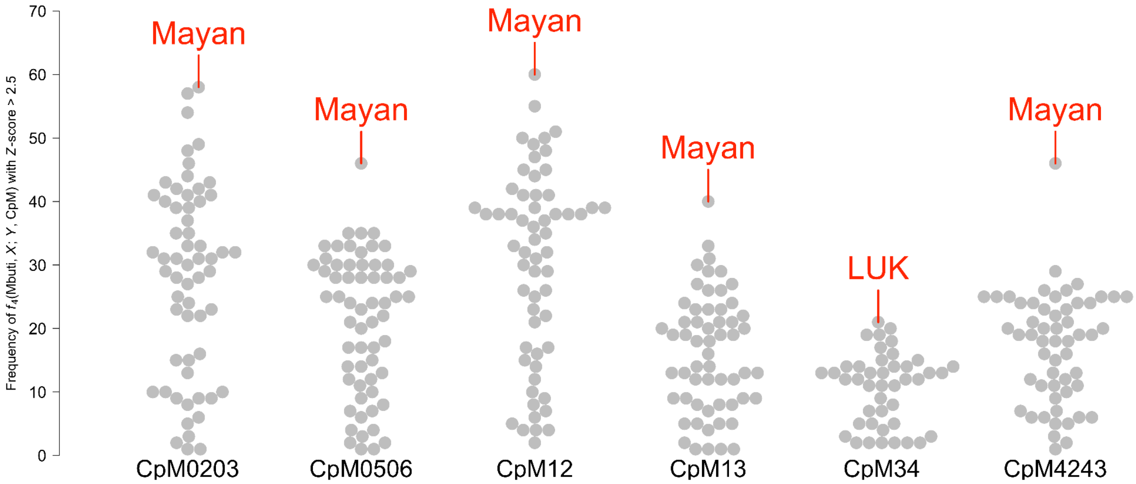


**Fig. S7. Distributions of populations ranked by significant genetic affinities in $f_4$(Mbuti, *X*; *Y*, CpM).** The plots illustrate the frequency with which a population stands out in the $f_4$-tests. Each plot represents a population *X* who exhibits an excess genetic affinity to the Copan individuals compared to a population *Y*, with a *Z*-score > 2.5. The present-day Maya population emerges as the most prevalent outlier in five out of six Copan individuals. Detailed rankings of the top five populations in each Classic Copan individual can be found in table S2.

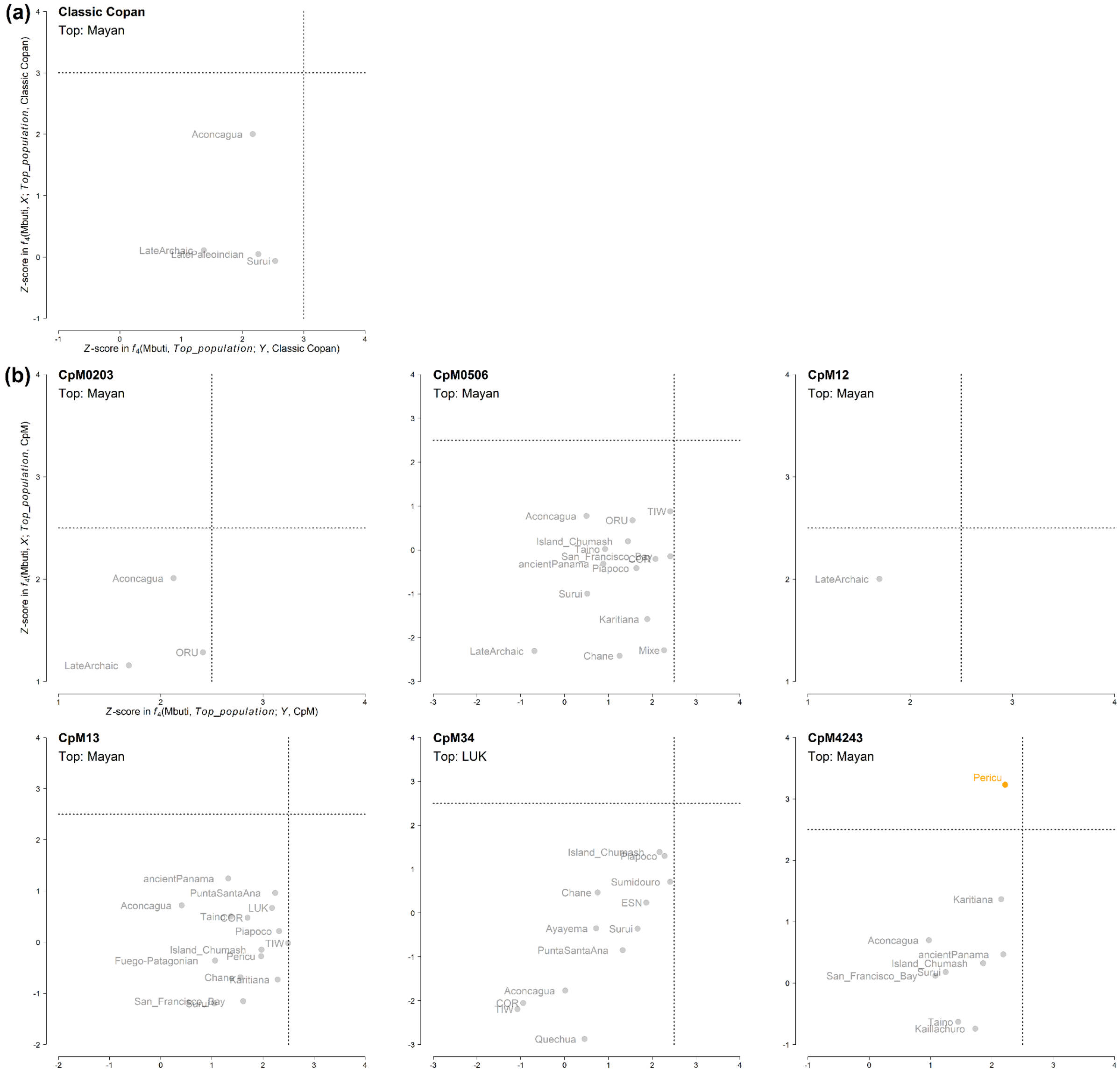


**Fig. S8. Classic Copan at (a) the population or (b) the individual level with *Z*-scores based on two different forms of *f*4-tests.** The *x*-axis illustrates *Z*-scores derived from *f*4(Mbuti, *Top_population*; *Y*, Classic Copan/CpM), while the *y*-axis displays those from *f*4(Mbuti, *X*; *Top_population*, Classic Copan/CpM). Top is a population ranked as the top in the *f*4(Mbuti, *X*; *Y*, Classic Copan/CpM) tests as shown in Figure 2. None of the populations that do not support genetic affinity between the top populations and the Copan (see the *Z*-scores on the *x*-axis, which are all below statistical significance) demonstrates a stronger genetic link to the Copan compared with the top populations (see the *y*-axis, where there is no population ), except for Pericu in CpM4243.

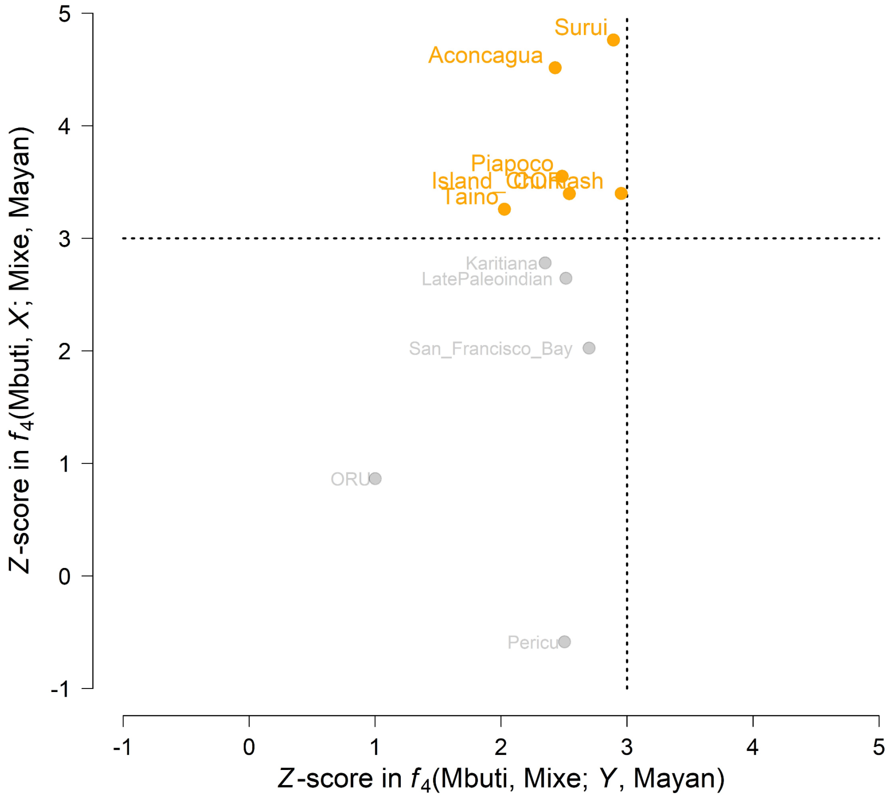


**Fig. S9. Populations with *Z*-scores based on two different forms of $f_4$-tests.** The *x*-axis illustrates *Z*-scores derived from $f_4$(Mbuti, Mixe; *Y*, Maya), while the *y*-axis displays those from $f_4$(Mbuti, *X*; Mixe, Maya). None of the populations included in the plot supports a significant genetic affinity between Mixe and Maya (see the *x*-axis), suggesting that they may share more affinity with Maya than Mixe, even though the genetic links between Mixe and Maya are evident in Figure 3. Indeed, some of these populations, which are Surui, Aconcagua, Piapoco, Island_Chumash, COR, and Taino, exhibit a stronger genetic affinity to Maya compared to Mixe (see the *y*-axis). These results indicate a potential dual ancestry of the Maya population (see Figure 3 and table S3).

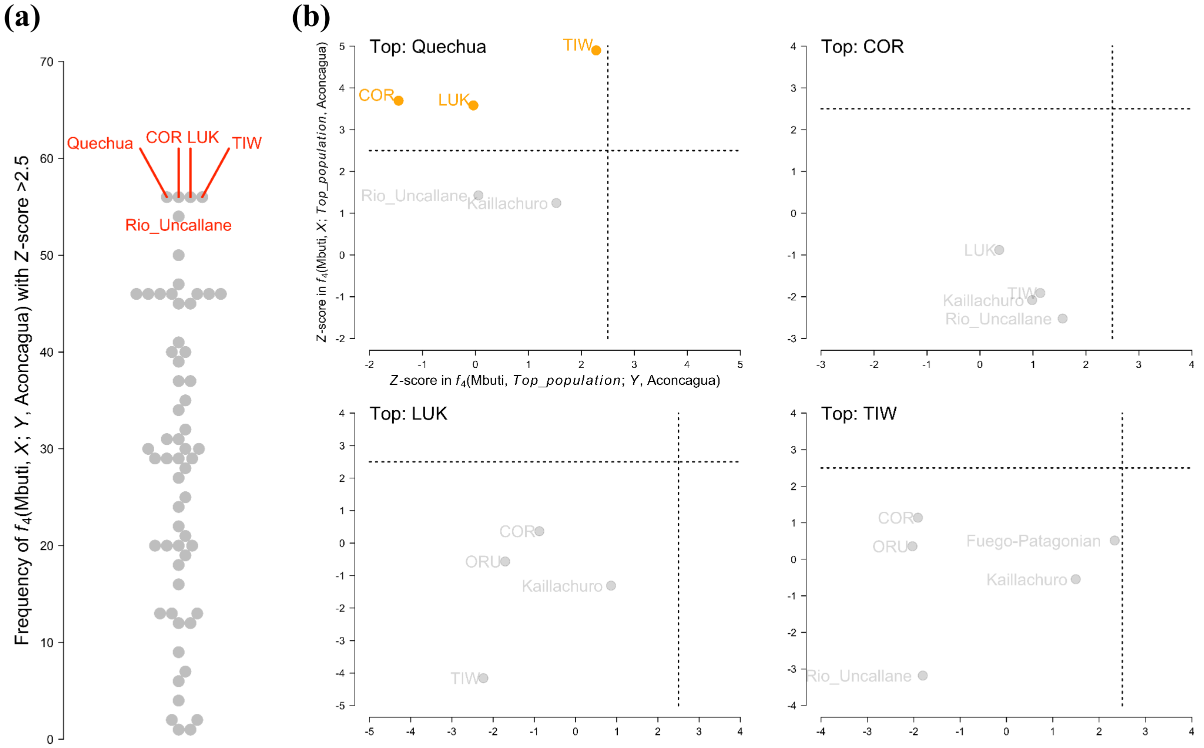


**Fig. S10. Exploring the genetic ancestry of Aconcagua. (a)** Genetic affinity of ancient and modern populations from Siberia and the Americas to Aconcagua is ranked by how many times they stand out as population *X* with *Z*-score > 2.5 in $f_4$(Mbuti, *X*; *Y*, Aconcagua). There are four populations with the highest occurrence as population *X*: Quechua, COR, LUK, and TIW. The second from the top populations is Rio_Uncallan. All of these populations are genetically grouped as ancient and modern populations sharing South American Farmer (SAF) ancestry (see ADMIXTURE results in fig. S6). **(b)** Populations with *Z*-scores based on two different forms of $f_4$-tests. The *x*-axis illustrates *Z*-scores derived from $f_4$(Mbuti, *Top_population*; *Y*, Aconcagua), while the *y*-axis displays those from $f_4$(Mbuti, *X*; *Top_population*, Aconcagua). Top is a population ranked as the top in the $f_4$(Mbuti, *X*; *Y*, Aconcagua) tests as shown in (a). Most of the populations that do not support genetic affinity between the top populations and Aconcagua (see the *Z*-scores on the *x*-axis, which are all below statistical significance) show no extra genetic affinity to Aconcagua compared with the top populations (see the *y*-axis, where there is no population ), except for COR, TIW, and LUK in the case of Quechua. Given that COR, TIW, and LUK are among the top populations all sharing SAF ancestry, these results confirm their strong genetic affinities to Aconcagua, rather than indicating dual ancestry of Quechua and these populations.

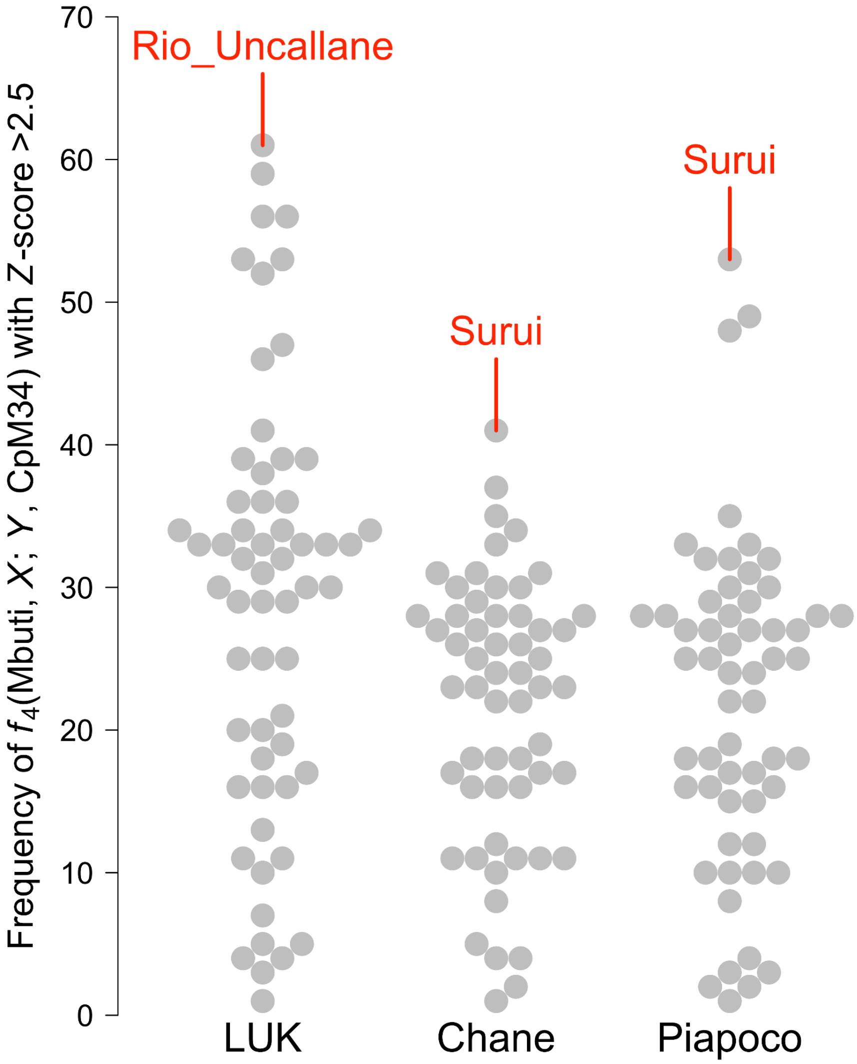


**Fig. S11. Rankings of populations in their significant genetic affinities to LUK, Chane, and Piapoco.** The plots illustrate the frequency with which a population stands out in the $f_4$-tests, with the form of $f_4$(Mbuti, $X$; $Y$, LUK/Chane/Piapoco). LUK, Chane, and Piapoco are top three populations in their genetic affinities to CpM34, as shown in table S2. The South American Farmers, Rio_Uncallane, emerges as the top population in the genetic affinities to LUK, while both of Chane and Piapoco support their robust genetic affinities to Surui, representing Amazonian ancestry (see PCA and ADMIXTURE results in figs. S4 and S6).

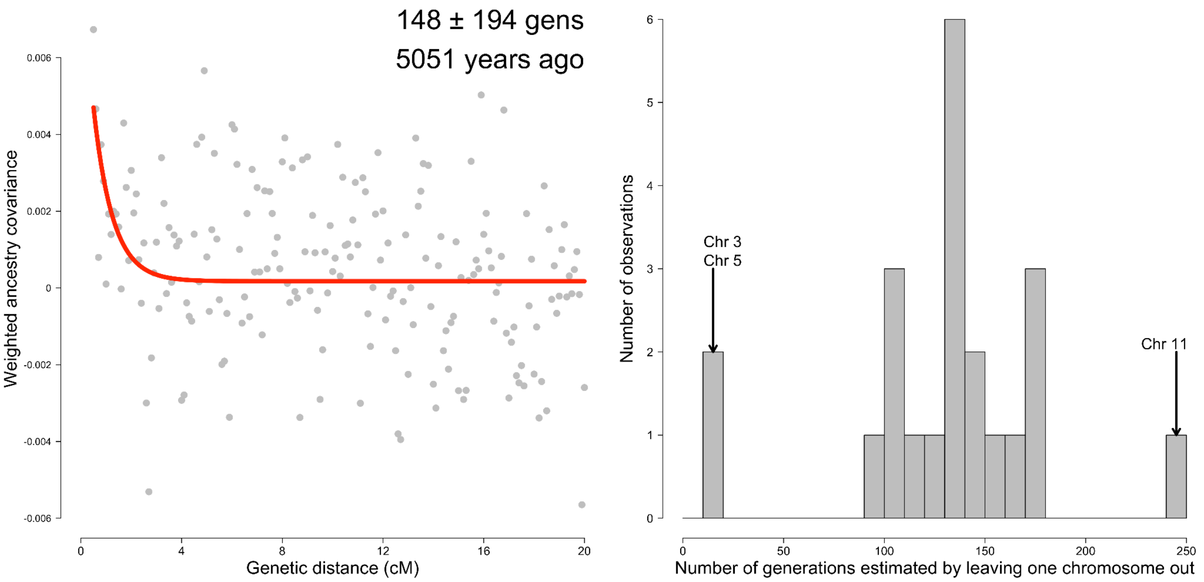


**Fig. S12. Dating admixture in the Classic Copan by the *DATES* program**. **Left:** The plot shows the exponential decay of weighted ancestry covariance ($y$-axis) with genetic distance ($x$-axis), in which a decay rate depends on the time since admixture. Fitting starts at a genetic distance of 0.45 centimorgan (cM). The estimates are converted to a number of years before present by adding the values to the mean age of samples (*i.e.*, ), with an assumption of 25 years per generation. **Right:** Due to large standard errors computed from a weighted block jackknife approach, the estimated dates obtained by leaving one autosomal chromosome out in each run are plotted as a histogram. The majority of the dates fall within a range of 90 to 180 generations, with notable outliers observed for Chromosomes 3, 5, and 11.

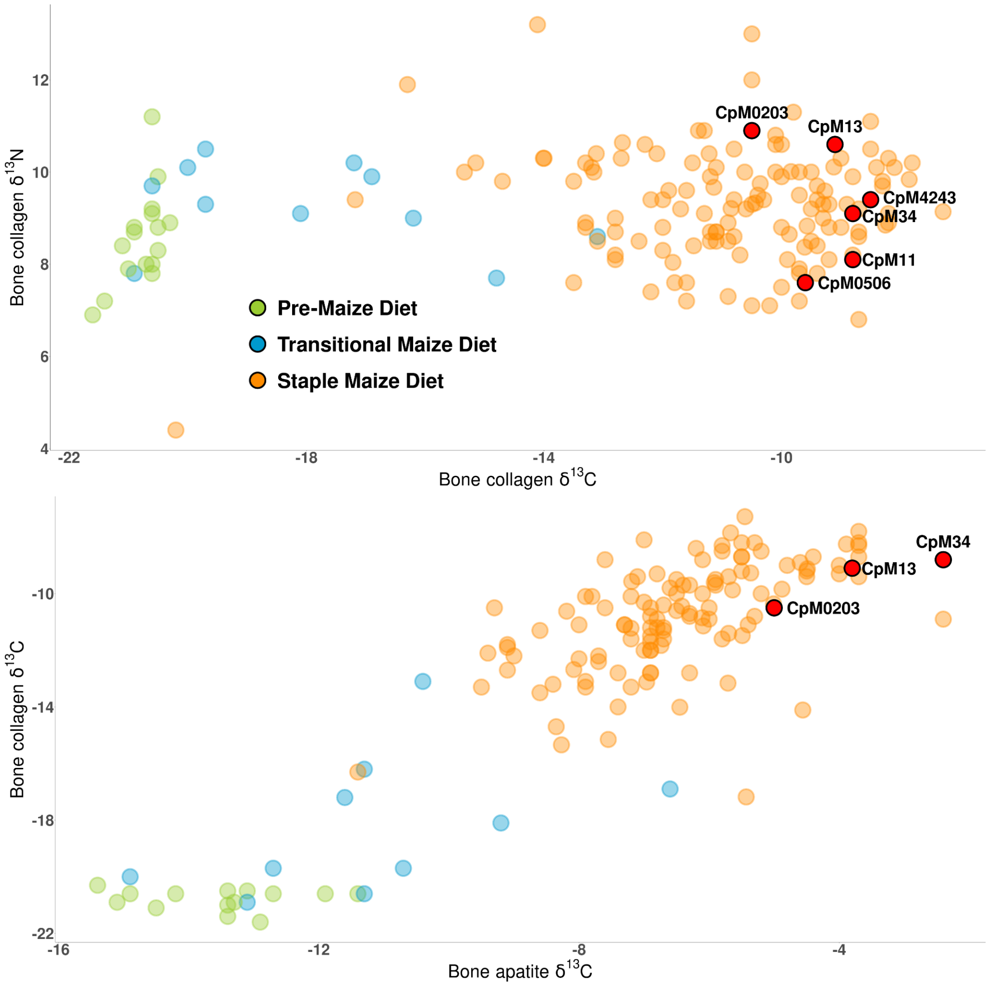


**Fig. S13. Isotopic profiles of the Classic Copan individuals (red) and ancient Maya individuals with pre-maize (green), transitional maize (blue), and stape maize diets (orange).** The top and bottom plots show values of $\delta^{13}C_{collagen}$ (*x*-axis) and $\delta^{13}N_{collagen}$ (*y*-axis) or those of $\delta^{13}C_{apatite}$ (*x*-axis) and $\delta^{13}C_{collagen}$ (*y*-axis), respectively. Due to a lack of isotopic data, CpM12 or CpM0506, CpM11, and CpM4243 are not included in the top or bottom plot. All data, except for the Classic Copan individuals, are referred from previous studies (*25*, *38*).

***Table S1. A summary of all published data included in the study.***

| Population group | Individual ID | Median calBP[a] | Removed as 1st degree relatives | #Inds included | Downloaded data format | Data type | References |
|---|---|---|---|---|---|---|---|
| AG2 | AG2 | 16,913 | | 1 | BAM | WGS | (*64*) |
| MA1 | MA1 | 24,157 | | 1 | BAM | WGS | |
| Birnirk | A2_N79, A3_N181 | 1,327-*NA* | | 2 | FASTQ | WGS | (*74*) |
| Pre-Dorset | XIV_H168 | 3,924 | | 1 | FASTQ | WGS | |
| Middle_Dorset | NiNg_1_A6, MARC480, MARC481, MARC483, MARC484, MARC485, MARC486, MARC487, MARC488, MARC1490, MARC1491, MARC1493, XIV_C3_4, Alarnerk3 | 1,921-1,162 | | 14 | FASTQ | WGS | |
| Late_Dorset | XIV_H126, XIV_C340 | 1,324-765 | | 2 | FASTQ | WGS | |
| Norton | B_003_1 | *NA* | | 1 | FASTQ | WGS | |
| Saqqaq | Qt1, L13 | 4,706-3,677 | | 2 | FASTQ | WGS | |
| Thule | XIVC_748, KAL1245, KAL1422, XIVC_746 | 370-357 | | 4 | FASTQ | WGS | |
| Clovis | Anzick_1 | 12,632 | | 1 | FASTQ | WGS | (*67*) |
| Fuego-Patagonian | AM71, MA572, MA575, MA577, 890, 894, 895 | ~200* | | 7 | FASTQ | WGS | (*75*) |
| LucyIslands | 939 | 6,075 | | 1 | FASTQ | WGS | |
| MummyMaderasEncoC2 | Chinchorro | 5,839 | | 1 | FASTQ | WGS | |
| OldMissionPoint | MARC1492 | 387 | | 1 | FASTQ | WGS | |
| Paleoamericans | BC23, BC25, BC27, BC28, BC29, BC30 | 800*-300* | | 6 | FASTQ | WGS | |
| Pre-Columbian | F9, MOM6 | >500* | | 2 | FASTQ | WGS | |
| SerradaCapivara | Enoque65 | 3,559 | | 1 | FASTQ | WGS | |
| Kennewick | Kennewick | 8,545 | | 1 | FASTQ | WGS | (*76*) |
| Kaillachuro | K1 | ~3,800* | | 1 | FASTQ | WGS | (*28*) |
| Rio_Uncallane | IL2, IL3, IL4, IL5, IL7 | 1,514-1,806 | | 5 | FASTQ | WGS | |
| SMP | SMP5 | 6,822 | | 1 | FASTQ | WGS | |
| USR1 | USR1 | 11,435* | Possible first cousins or half-siblings | 1 | FASTQ | WGS | (*65*) |
| USR2 | USR2 | | | 1 | FASTQ | WGS | |
| Aconcagua | Aconcagua | ~500* | | 1 | FASTQ | WGS | (*29*) |
| Ayayema | A460 | 5,121 | | 1 | FASTQ | WGS | |
| BigBar | 19651 | 5,664 | | 1 | FASTQ | WGS | |
| Lovelock | Lovelock1, Lovelock2, Lovelock3, Lovelock4 | 1,950-667 | | 4 | FASTQ | WGS | |
| PuntaSantaAna | SA5832 | 7,293 | | 1 | FASTQ | WGS | |
| SpiritCave | AHUR_2064 | 10,946 | | 1 | FASTQ | WGS | |

| | | | | | | | |
|---|---|---|---|---|---|---|---|
| Sumidouro | Sumidouro4, Sumidouro5, Sumidouro6, Sumidouro7, Sumidouro8 | 10,408-9,806 | | 5 | FASTQ | WGS | |
| TrailCreek | TrailCreek | 9,020 | | 1 | FASTQ | WGS | |
| Alaskan_Athabaskan | 523a | *NA* | | 1 | BAM | WGS | |
| ASO | CK01, CK02, CK03_merged, CK04, CK07_merged, CK08, CK09_merged, CK10_merged, CK13_merged, LU01, LU02, LU03, LU04, LU05, LU06 | 4,845-397 | | 15 | BAM | WGS | |
| ASO_Huron-Wendat | RM83, RM85 | 500[**] | | 2 | BAM | WGS | |
| Baja_Mexico | B04, MX01 | *NA* | | 2 | BAM | WGS | |
| Island_Chumash | CR01_merged, SM01, SM02_merged | 663-122 | | 3 | BAM | WGS | |
| LSCI | CT01_merged, CT02, SC01, SC02_merged, SC03_merged, SC04, SC05_merged, SC06, SC07 | 661-70 | | 9 | BAM | WGS | |
| Mainland_Chumash | CH01, NC, PS02_merged, PS03_merged, PS04, PS06_merged, PS07_merged, PS09_merged, PS12, PS13, PS17_merged, PS18_merged, PS22, PS23, PS26_merged, PS30, PS34 | 1,780-1,335 | | 17 | BAM | WGS | (*66*) |
| Pericu | B03 | *NA* | | 1 | BAM | WGS | |
| ESN | SN04, SN17_merged, SN20, SN25, SN31, SN32_merged, SN37, SN39, SN40, SN41, SN44_merged, SN45, SN48, SN54, SN55, SN56, SN57, SN58, SN59, SN60_merged | 4,635-3,691 | | 20 | BAM | WGS | |
| LSN | SN01, SN03, SN09, SN10, SN11_merged, SN12, SN13_merged, SN15, SN16, SN38, SN43, SN50, SN51, SN52, SN53 | 2,045-281 | | 15 | BAM | WGS | |
| San_Francisco_Bay | Ala1 | *NA* | | 1 | BAM | WGS | |
| Taino | Taino9606 | 1,066 | | 1 | FASTQ | WGS | (*68*) |
| Coropuna (COR) | CO001, CO066, CO154, CO193 | 1,040-390 | | 4 | FASTQ | WGS | |
| Lukurmata (TIW) | TW013, TW020, TW027, TW028 | 1,665-480 | | 4 | FASTQ | WGS | |
| Oruro (ORU) | TW033 | 535 | | 1 | FASTQ | WGS | (*27*) |
| Tiwanaku (TIW) | TW004, TW008, TW013, TW033, TW056, TW060, TW061, TW063, TW097 | 1,225-865 | | 8 | FASTQ | WGS | |
| ancientPanama | PAPV_114_LTP, PAPV_117_RTP, PAPV_118_RTP, PAPV_146_TP, PAPV_167_RTP, PAPV_172_LTP, PAPV_173_RTP, | 1,317-572 | | 9 | FASTQ | WGS | (*26*) |

| | | | | | | | |
|---|---|---|---|---|---|---|---|
| | PAPV_174_LTP, PAPV_175_LTP | | | | | | |
| LateArchaic | I24540, I13267, I5454, I3442, I19942, I7544, I24542, I19950, I6236, I5455, I8041, I7543 | 5,570**-3,850 | | 12 | FASTQ | Capture | (*25*) |
| LatePaleoindian | I13268, I19169 | 9,540-8,785 | I19170 removed as 1st degree to I19169 | 2 | FASTQ | Capture | |

[a]The values show the oldest and most recent ages if multiple samples were sequenced from the same site. BP stands for "Before Present" where "present" refers to the year 1950. *NA* stands for "Not Available" data.
*Direct datation from the sample was unavailable, so the date was estimated based on archeological context or charcoal from the archeological site.
**Age is not calibrated (not adjusted for the atmospheric fluctuations of $C^{14}$).

***Table S2. Top five populations with Z-score >3.0 or 2.5 in the form of $f_4$(Mbuti, X; Y, Classic Copan/CpM).***

| **Rank** | **Population level** | **Individual level** | | | | | |
|---|---|---|---|---|---|---|---|
| | **Classic Copan** | **CpM0203** | **CpM0506** | **CpM12** | **CpM13** | **CpM34** | **CpM4243** |
| **1** | Mayan | Mayan | Mayan | Mayan | Mayan | LUK | Mayan |
| **2** | ancientPanama | ancientPanama | Zapotec<br>Piapoco<br>ancientPanama | Piapoco | Piapoco | Chane | Karitiana |
| **3** | Mixe | Mixe | Mainland_Chumash<br>LSN<br>LSCI<br>Surui<br>Aconcagua | LSCI | ancientPanama | Karitiana<br>Piapoco | Surui |
| **4** | Piapoco | LUK | TIW | LSN<br>ancientPanama | Mixe | ancientPanama | Island_Chumash<br>Mixe |
| **5** | LateArchaic | LateArchaic | LUK | Karitiana | Chane<br>Taino | Mayan | Zapotec<br>Rio_Uncallane<br>Quechua<br>LUK<br>LSCI<br>ancientPanama<br>Aconcagua |

***Table S3. Testing fittings of two-way admixture models between Mixe and each of potential sources to the genetic ancestry of Mayan by qpAdm.***

| **Model parameters (Target: Maya)** | | **Source 2** | | | | | |
|---|---|---|---|---|---|---|---|
| | | **Aconcagua** | **COR** | **Island Chumash** | **Piapoco** | **Surui** | **South American Farmer** |
| **Model** | | **Two-way admixture** | | | | | |
| **DoF** | | 11 | 11 | 11 | 11 | 11 | 11 |
| $x^2$ | | 8.365 | 7.499 | 11.893 | 9.918 | 12.053 | 7.729 |
| **Model** | | **Single ancestry with Mixe** | | | | | |
| **DoF** | | 12 | 12 | 12 | 12 | 12 | 12 |
| $x^2$ | | 14.824 | 14.806 | 14.843 | 14.854 | 14.839 | 14.858 |
| **Model** | | **Single ancestry with Source 2** | | | | | |
| **DoF** | | 12 | 12 | 12 | 12 | 12 | 12 |
| $x^2$ | | 52.698 | 75.296 | 41.644 | 67.557 | 157.96 | 76.641 |
| **Model selection** | | **Testing dual ancestry vs. single ancestry scenarios** | | | | | |
| **Nested *p*-value against Mixe** | | 0.011 | 0.007 | 0.086 | 0.026 | 0.095 | 0.008 |
| **Nested *p*-value against Source 2** | | 0.000 | 0.000 | 0.000 | 0.000 | 0.000 | 0.000 |
| **Best fitting model** | | Admixture | Admixture | Mixe-only | Admixture | Mixe-only | Admixture |
| **Admixture proportions** | **Mixe** | 75.5% ± 9.5% | 71.7% ± 9.6% | - | 79.0% ± 10.0% | - | 75.5% ± 8.7% |
| | **Source 2** | 24.5% ± 9.5% | 28.3% ± 9.6% | - | 21.0% ± 10.0% | - | 24.5% ± 8.7% |

***Table S4. Testing fittings of two-way admixture models between Quechua and each of potential sources to the genetic ancestry of Aconcagua by qpAdm.***

| **Model parameters (Target: Aconcagua)** | **Source 2** | | |
|---|---|---|---|
| | **COR** | **LUK** | **TIW** |
| **Model** | **Two-way admixture** | | |
| **DoF** | 11 | 11 | 11 |
| $x^2$ | 10.672 | 7.532 | 7.462 |
| **Model** | **Single ancestry with Quechua** | | |
| **DoF** | 12 | 12 | 12 |
| $x^2$ | 33.207 | 33.226 | 33.233 |
| **Model** | **Single ancestry with Source 2** | | |
| **DoF** | 12 | 12 | 12 |
| $x^2$ | 10.626 | 12.16 | 7.511 |
| **Model selection** | **Testing dual ancestry vs. single ancestry scenarios** | | |
| **Nested *p*-value against Quechua** | 0.000 | 0.000 | 0.000 |
| **Nested *p*-value against Source 2** | 1.000 | 0.031 | 0.825 |
| **Best fitting model** | COR-only | Admixture | TIW-only |
| **Quechua** | - | *NA* | - |
| **Source** | - | *NA* | - |

***Table S5. Testing fittings of admixture models between Mixe and Late Archaic ancestry or between Mixe and South America Farmer (SAF) ancestry to genetic ancestry of Mayan by qpAdm.***

| **Modelling parameters** | **Different admixture scenarios** | | | | | | |
|---|---|---|---|---|---|---|---|
| | **Three-way admixture** | **Two-way admixture** | | | **Single ancestry** | | |
| **DoF** | 10 | 11 | 11 | 11 | 12 | 12 | 12 |
| $\chi^2$ | 4.794 | 15.279 | 12.974 | 7.707 | 22.576 | 76.628 | 14.825 |
| **Tail probability** | 0.904 | 0.170 | 0.295 | 0.739 | 0.032 | 1.807e-11 | 0.251 |
| **Mixe** | -0.690 | - | 0.325 | 0.753 | - | - | 1.000 |
| **Late Archaic** | 2.081 | 1.071 | 0.675 | - | 1.000 | - | - |
| **SAF** | -0.391 | -0.071 | - | 0.247 | - | 1.000 | - |
| **Model fit** | -[a] | -[a] | Good fit | Good fit | -[b] | -[b] | Good fit |
| **Nested *p*-value** | - | - | Mixe<br>0.174<br>Late Archaic<br>1.443e-15 | Mixe<br>0.008<br>SAF<br>1.110e-16 | - | | |

[a]The admixture is not supported from modelling due to negative values of admixture coefficients.
[b]The admixture is not supported from modelling due to tail probabilities below 0.05.

***Table S6. Testing fittings of two-way admixture models to genetic ancestry of Classic Copan or LateArchaic by qpAdm.***

<table>
<tr><th rowspan="2">Target population</th><th colspan="5">Two-way admixture</th><th colspan="4">Single ancestry models</th></tr>
<tr><th>DoF</th><th>$x^2$</th><th>Tail prob</th><th>Mesoamerican</th><th>SAF</th><th>DoF</th><th>$x^2$</th><th>Tail prob</th><th>Nested p-value</th></tr>
<tr><td rowspan="4">Classic Copan</td><td rowspan="4">11</td><td rowspan="4">12.068</td><td rowspan="4">0.359</td><td rowspan="4">21.0% ± 8.4%</td><td rowspan="4">79.0% ± 8.4%</td><td colspan="4">Mesoamerican-only</td></tr>
<tr><td>12</td><td>75.672</td><td>2.742e-11</td><td>0.000</td></tr>
<tr><td colspan="4">SAF-only</td></tr>
<tr><td>12</td><td>18.515</td><td>0.101</td><td>0.011</td></tr>
<tr><td rowspan="4">LateArchaic</td><td rowspan="4">11</td><td rowspan="4">16.714</td><td rowspan="4">0.117</td><td rowspan="4">8.1% ± 16.0%</td><td rowspan="4">91.9% ± 16.0%</td><td colspan="4">Mesoamerican-only</td></tr>
<tr><td>12</td><td>71.346</td><td>1.792e-10</td><td>0.000</td></tr>
<tr><td colspan="4">SAF-only</td></tr>
<tr><td>12</td><td>17.006</td><td>0.149</td><td>0.589</td></tr>
</table>

***Table S7. Results for f4-statistics with the form of f4(Mbuti, X; Mayan, Classic Copan).***

| Pop1 | Pop2 (*X*) | Pop3 | Pop4 | $f_4$ | sd | *Z*-score (bold if *Z* > 3.0) |
|---|---|---|---|---|---|---|
| Mbuti | **LSCI** | Mayan | Classic Copan | 0.001042 | 0.000222 | **4.692** |
| | **LSN** | | | 0.000895 | 0.000211 | **4.239** |
| | **TIW** | | | 0.000798 | 0.000192 | **4.164** |
| | **Rio_Uncallane** | | | 0.000760 | 0.000217 | **3.496** |
| | **Sumidouro** | | | 0.001011 | 0.000317 | **3.188** |
| | **Mainland_Chumash** | | | 0.000722 | 0.000227 | **3.185** |
| | **Kaillachuro** | | | 0.001021 | 0.000337 | **3.031** |
| | ancientPanama | | | 0.000612 | 0.000214 | 2.866 |
| | LUK | | | 0.000652 | 0.000232 | 2.814 |
| | Baja_Mexico | | | 0.000886 | 0.000326 | 2.720 |
| | COR | | | 0.000596 | 0.000219 | 2.718 |
| | ORU | | | 0.000919 | 0.000374 | 2.459 |
| | Fuego-Patagonian | | | 0.000650 | 0.000278 | 2.343 |
| | SMP | | | 0.000990 | 0.000434 | 2.281 |
| | Pericu | | | 0.000734 | 0.000335 | 2.192 |
| | Lovelock | | | 0.000475 | 0.000224 | 2.124 |
| | ASO | | | 0.000566 | 0.000273 | 2.077 |
| | ESN | | | 0.000497 | 0.000244 | 2.039 |
| | Aconcagua | | | 0.000609 | 0.000304 | 2.004 |
| | Ayayema | | | 0.000573 | 0.000302 | 1.897 |
| | Island_Chumash | | | 0.000460 | 0.000251 | 1.829 |
| | Taino | | | 0.000470 | 0.000278 | 1.69 |
| | PuntaSantaAna | | | 0.000504 | 0.000311 | 1.618 |
| | Colonist | | | 0.000427 | 0.000284 | 1.501 |
| | Alaskan_Athabaskan | | | 0.000379 | 0.000275 | 1.382 |
| | BigBar | | | 0.000398 | 0.000288 | 1.379 |
| | Pre-Dorset | | | 0.001673 | 0.001271 | 1.316 |
| | TrailCreek | | | 0.000401 | 0.000309 | 1.300 |
| | Piapoco | | | 0.000289 | 0.000228 | 1.268 |
| | SpiritCave | | | 0.000388 | 0.000309 | 1.256 |
| | USR1 | | | 0.000325 | 0.000279 | 1.166 |
| | Birnirk | | | 0.001821 | 0.001661 | 1.096 |
| | Tlingit | | | 0.000203 | 0.000194 | 1.047 |
| | Karitiana | | | 0.000252 | 0.000251 | 1.003 |

| | | | | | | |
|---|---|---|---|---|---|---|
| | San_Francisco_Bay | | | 0.000328 | 0.000342 | 0.960 |
| | Middle_Dorset | | | 0.000362 | 0.000428 | 0.846 |
| | MA1 | | | 0.000215 | 0.000267 | 0.803 |
| | Anzick | | | 0.000229 | 0.000296 | 0.774 |
| | AG2 | | | 0.000243 | 0.000323 | 0.752 |
| | Norton | | | 0.001401 | 0.001870 | 0.749 |
| | colonialPanama | | | 0.000588 | 0.001047 | 0.562 |
| | USR2 | | | 0.000222 | 0.000405 | 0.548 |
| | ASO_4000 | | | 0.000120 | 0.000294 | 0.410 |
| | MummyMaderasEncoC2 | | | 0.004735 | 0.013936 | 0.340 |
| | Pre-Columbian | | | 0.000283 | 0.000918 | 0.308 |
| | ASO_Huron-Wendat | | | 0.000086 | 0.000315 | 0.273 |
| | Kennewick | | | 0.000103 | 0.000409 | 0.252 |
| | Saqqaq | | | 0.000230 | 0.001172 | 0.196 |
| | Paleoamericans | | | 0.000065 | 0.000410 | 0.158 |
| | LateArchaic | | | 0.000057 | 0.000517 | 0.111 |
| | LatePaleoindian | | | 0.000062 | 0.001287 | 0.048 |
| | Surui | | | -1.6E-05 | 0.000256 | -0.063 |
| | LucyIslands | | | -0.00005 | 0.000339 | -0.148 |
| | Quechua | | | -3.9E-05 | 0.000222 | -0.175 |
| | SerradaCapivara | | | -0.00130 | 0.005783 | -0.225 |
| | Thule | | | -0.00044 | 0.001587 | -0.276 |
| | OldMissionPoint | | | -0.00024 | 0.000510 | -0.462 |
| | Late_Dorset | | | -0.00023 | 0.000353 | -0.661 |
| | Zapotec | | | -0.00017 | 0.000240 | -0.706 |
| | Hawaiian | | | -0.00017 | 0.000218 | -0.789 |
| | Mixe | | | -0.00020 | 0.000222 | -0.905 |
| | Pima | | | -0.00033 | 0.000248 | -1.336 |
| | Mixtec | | | -0.00030 | 0.000212 | -1.414 |
| | Chane | | | -0.00043 | 0.000285 | -1.512 |

***Table S8. Testing fittings of two-way admixture models to genetic ancestry of Classic Copan individuals by qpAdm.***

| **Model parameters** | **Classic Copan individuals (each Copan individual is used as a target for admixture modelling)** | | | | | |
|---|---|---|---|---|---|---|
| | **CpM0203** | **CpM0506** | **CpM12** | **CpM13** | **CpM34** | **CpM4243** |
| **Model** | **Two-way admixture** | | | | | |
| **DoF** | 11 | 11 | 11 | 11 | 11 | 11 |
| $x^2$ | 9.431 | 17.804 | 7.508 | 10.581 | 16.539 | 8.069 |
| **Tail prob** | 0.582 | 0.086 | 0.757 | 0.479 | 0.122 | 0.707 |
| **Admixture proportions** | **Source 1** | | | | | |
| | Mesoamerican | Mesoamerican | Mesoamerican | Mesoamerican | Amazonian | Mesoamerican |
| | 5.8% ± 13.2% | 37.0% ± 15.1% | 11.5% ± 14.8% | 47.4% ± 18.6% | 26.5% ± 18.5% | 4.2% ± 19.2% |
| | **Source 2** | | | | | |
| | SAF (Aconcagua+COR) | SAF (Aconcagua+COR) | SAF (Aconcagua+COR) | SAF (Aconcagua+COR) | SAF (LUK) | SAF (Aconcagua+COR) |
| | 94.2% ± 13.2% | 63.0% ± 15.1% | 88.5% ± 14.8% | 52.6% ± 18.6% | 73.5% ± 18.5% | 95.8% ± 19.2% |
| **Model** | **Single ancestry with Source 1** | | | | | |
| **DoF** | 12 | 12 | 12 | 12 | 12 | 12 |
| $x^2$ | 54.258 | 34.599 | 43.692 | 19.614 | 34.340 | 38.929 |
| **Tail prob** | 2.458e-07 | 5.423e-04 | 1.723e-05 | 0.075 | 5.960-e04 | 1.081e-04 |
| **Model** | **Single ancestry with Source 2** | | | | | |
| **DoF** | 12 | 12 | 12 | 12 | 12 | 12 |
| $x^2$ | 9.608 | 25.958 | 8.195 | 18.577 | 19.994 | 8.098 |
| **Tail prob** | 0.650 | 0.011 | 0.770 | 0.992 | 0.067 | 0.777 |
| **Model selection** | **Testing dual ancestry vs. single ancestry scenarios** | | | | | |
| **Nested *p*-value against Source 1** | 0.000 | 0.000 | 0.000 | 0.003 | 0.000 | 0.000 |
| **Nested *p*-value against Source 2** | 0.674 | 0.004 | 0.407 | 0.005 | 0.063 | 0.865 |
